# Efficient numerosity estimation under limited time

**DOI:** 10.1101/2023.07.03.547493

**Authors:** Joseph A. Heng, Michael Woodford, Rafael Polania

## Abstract

The ability to rapidly estimate non-symbolic numerical quantities is a well-conserved sense across species with clear evolutionary advantages. Despite its importance, the rapid representation and estimation of numerosity is surprisingly imprecise and biased. However, a formal explanation for this seemingly irrational behavior remains unclear. We develop a unified normative theory of numerosity estimation that parsimoniously incorporates in a single framework information processing constraints alongside Brownian diffusion noise to capture the effects of time exposure of sensory estimations, logarithmic encoding of numerosity representations, and optimal inference via Bayesian decoding. We show that for a given allowable biological capacity constraint our model naturally endogenizes time perception during noisy efficient encoding to predict the complete posterior distribution of numerosity estimates. This model accurately predicts many features of human numerosity estimation as a function of temporal exposure, indicating that humans can rapidly and efficiently sample numerosity information over time. Additionally, we demonstrate how our model fundamentally differs from a thermodynamically-inspired formalization of bounded rationality, where information processing is modeled as acting to shift away from default states. The mechanism we propose is the likely origin of a variety of numerical cognition patterns observed in humans and other animals.

**Author summary:** Humans have the ability to estimate the number of elements in a set without counting. We share this ability with other species, suggesting that it is evolutionarily relevant. However, despite its relevance, this sense is variable and biased. What is the origin of these imprecisions? We take the view that they are the result of an optimal use of limited neural resources. Because of these limitations, stimuli are encoded with noise. The observer then optimally decodes these noisy representations, taking into account its knowledge of the distribution of stimuli. We build on this view and incorporate stimulus presentation time (or contrast) directly into the encoding process using Brownian motion. This model can parsimoniously predict key characteristics of our perception and outperforms quantitatively and qualitatively a popular modeling approach that considers resource limitations at the stage of the response rather than the encoding.

## Introduction

The ability to rapidly represent and estimate non-symbolic numerical quantities is a fundamental cognitive function for behavior in humans and other animals, which may have emerged during evolution to support fitness maximization [1]. Since the properties of numerosity estimation started to be studied nearly a century ago, it has been commonly observed that the representation and estimation of numerical quantities are imprecise and biased [2]. Despite the importance of numerosity estimation for various cognitive processes and ultimately survival, the questions remain: what are the origins of the observed variability and biases in numerosity estimations? Are these deviations efficient and predictable when organisms are urged to rapidly estimate numerical quantities?

Extensive empirical research in the representation and estimation of non-symbolic numerical quantities has consistently reported and studied various features that characteristically emerge during numerosity estimation, including: (i) subitizing small numbers [3]; (ii) overestimation of small numbers (outside the subitization range) and underestimation of large numbers [4], with especially biased estimates in the case of larger numbers [5]; (iii) a coefficient of variation that is approximately constant across all numerosities, a property termed scalar variability [6]; and (iv) estimation acuity modulated by duration of stimulus presentation and sensory reliability [7]. But do all the above-mentioned behavioral patterns have a common origin?

The last decades have been marked by the development of models of behavior in which perception has been proposed to be instantiated as a Bayesian inference process. This suggests that our nervous system jointly considers the environmental (or contextual) distribution of sensory stimuli and the unreliability of the signals perceived by the observer. This approach has been instrumental in explaining in a parsimonious manner a variety of behavioral biases including underestimation, overestimation, and the degree of variability of estimated magnitudes and quantities [8]. However, this approach does not explicitly consider the different sets of constraints that biological systems face when interacting with the environment. This is a fundamental aspect to consider in any formulation that attempts to explain the behavior of biological systems given the fact that organisms do not have unlimited biological resources or unlimited time to process sensory information from the environment, and moreover, neural computations are metabolically expensive [9]. Thus, it has been suggested that the observed variability and biases in our estimations of our sensory world emerge from fundamental principles of acquiring information from environmental regularities that should ultimately lead to developing efficient behavioral strategies [10–14].

Here we argue that many of the above-mentioned behavioral features emerging during numerosity estimation have a common origin: given biological constraints on information acquisition, numerosity estimation emerges from a system that efficiently considers, first, prior knowledge of the environment, second, information of the current numerosity being evaluated, and third, the amount of time available to process such information.

We develop a unified normative model of numerosity estimation that parsimoniously incorporates information constraints together with long modeling traditions of human and animal psychophysical performance in psychology and neuroscience: (i) Brownian diffusion noise to capture the effects of time exposure of sensory information [15], (ii) logarithmic encoding of numerosity representations [16], and (iii) optimal Bayesian decoding. As a result, we show that for a given allowable biological capacity constraint, our model naturally incorporates time perception during noisy efficient encoding to predict the corresponding posterior distribution of numerosity estimates via optimal Bayesian decoding. Here we refer to our approach as the “sequential-encoding/Bayesian-decoding” model, henceforth SEB.

We also consider a second well-known approach for studying bounded rationality inspired by principles of thermodynamics and statistical physics. This family of models assumes that given a default state (e.g., a default distribution over possible responses) and a sensory stimulus, the observer acts in a way such that they attempt to shift from the default state to a new state that matches as closely as possible the value of the sensory stimulus. Bounded rationality comes into play in the case of acting when only a given amount of change in information (energy invested) between the default and new state can be afforded. This class of models has been used in a wide range of applications [17–20], including recently to study how perceptual estimation under limited time relates to cognitive capacity and action responses [21]. Here we refer to this class of models as the “thermodynamically inspired model”, henceforth TIM.

A key contribution of our work is the formal demonstration that the two approaches that we consider here (SEB and TIM) are in fact classes of models with completely different views on bounded rationality. To avoid confusing them, we clarify their differences here. On the one hand, variability in the estimation responses in SEB is attributed to *sensing* costs, which generate noisy sensory encoding. On the other hand, in instantiations of TIM applied to sensory estimation, variability is generated by *acting* costs during response selection. Crucially, here we demonstrate that these two approaches applied to numerosity estimation lead to apparently similar but distinguishable quantitative and qualitative predictions that are identifiable and falsifiable. Our empirical tests applied to a large numerosity estimation data set provide a clear indication that humans follow a SEB rather than a TIM approach, meaning they can rapidly and efficiently sample numerosity information over time via an efficient noisy encoding and decoding process.

## Results

The presentation of our results is divided into three parts: First, we present our sequential-encoding/Bayesian-decoding model (SEB) which parsimoniously endogenizes perceptual exposure times in its likelihood function alongside parameters of the prior distribution for a given biological capacity bound. Second, we introduce the thermodynamically inspired model (TIM) applied to sensory estimation, and compare it with the SEB model. Third, we apply rigorous quantitative and qualitative model evaluations based on a large publicly available human numerosity estimation dataset (n=400 participants across four different experiments).

### A Bayesian model of numerosity estimation

Extensive behavioral and physiological work studying the representation of both non-symbolic and symbolic numerical quantities strongly suggests that internal representations *r* can be assumed to be encoded by a quantity that is proportional to the logarithm of the number *n* plus stimulus-independent random error that is assumed to be normally distributed and unbiased [5, 16, 22]

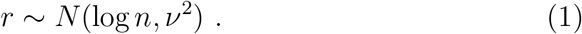

However, a key contribution of our work is to formally study how these perceptual errors may depend on stimulus duration *t* of the form

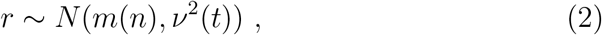

which represents the likelihood in our Bayesian framework. Here, *m*(*n*) is an affine function of *log*(*n*). Below, we will develop the theory by finding the parameters of the encoding process that minimize the MSE between the inputs and the estimation. That is, we aim to optimize the encoding function based on the variability of the representation which is a function of the sensory representation as a function of stimulus duration *t*.

We assume the prior distribution to be a log-normal distribution from which the true numerosity *n* is drawn to be

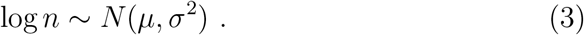

Note that *μ* and *σ* are the expected value and standard deviation of the random variable’s natural logarithm, and not the expectation and standard deviation of *n* itself.

While the distribution of various quantities in linguistics, economics, and ecology appears to be well-described by log-normal distributions [23], others have argued that power-law distributions approximately describe the empirical frequency of numbers in natural environments [24, 25]. We note, however, that the two-parameter family of possible log-normal prior distributions includes as a limiting case the power-law distributions (see S1 Appendix–Supplementary Note 1 for proof). In brief, we consider a normalized prior of the form

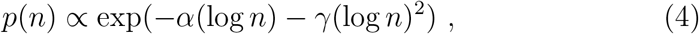

for some parameters *α, γ* with *γ* ≥ 0. If *γ >* 0, this corresponds to a log-normal prior, with *μ* = (1 − *α*)*/*(2*γ*), *σ*^2^ = 1*/*(2*γ*). If instead *γ* = 0 but *α >* 0, this corresponds to a power-law prior

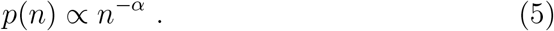

Thus, our model allows for the possibility that encoding and decoding are adapted to different priors that are learned for different contexts, rather than a single process being used in all contexts. However, in the following theoretical developments, we consider a log-normal prior for simplicity.

Here we assume that the objective of the decision-maker is to obtain numerosity estimates 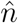 that minimize the MSE when stimuli are drawn from the prior distribution. It can be shown that this implies that conditional on *n*, the estimate 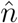 will be log-normally distributed (S1 Appendix–Supplementary Note 1)

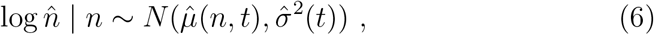

where 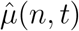 is an affine function of log *n*, and 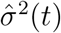 is independent of *n*. However, both 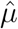 and 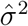 may depend on temporal numerosity processing *t*, as we formally elaborate below.

### Exposure time and precision of internal representations

A key hallmark in the development of our theoretical framework is that we now assume that the sensory evidence of the input stimulus is given by a Brownian motion with a drift that depends on the stimulus (formally defined below). Thus, by modeling sensory percepts in this way, we follow a long modeling tradition of process models of perception and action that includes the popular drift-diffusion model (DDM) [15]. Models of this kind have been used since the late 60s to account quantitatively for the way in which the accuracy of perceptual judgments is affected by manipulations of viewing time [26].

Formally, we now suppose that the internal representation *r* consists of the sample path of a Brownian motion *z*_*s*_ over a time interval 0 ≤ *s* ≤ *τ*, starting from an initial value *z*_0_ = 0. The drift-diffusion parameter *m* of the Brownian motion is assumed to depend on *n*, while its instantaneous variance *ω*^2^ is independent of *n*; the length of time *τ* for which the Brownian motion evolves is also independent of *n*, but depends on the viewing time *t*. In other words, we assume that the agent makes observations of momentary evidence 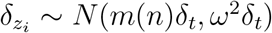 in small steps *i* for an infinitesimal duration *δ*_*t*_, where the accumulated evidence at step *k* is given by 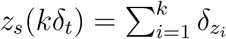.

Please note that the instantaneous variance *ω*^2^ of the diffusion process should not be confused with the resulting encoding noise *v*^2^ in the logarithmic space. In fact, our goal will be to formally find how *v* depends on elements that determine the diffusion process *m* and *ω* for given stimulus *n* and viewing time *t*.

More specifically, under the assumption that the particle position under Brownian motion is normally distributed with its parameters evolving as a function of *τ*, one can show that *r* is a draw from the distribution (S1 Appendix–Supplementary Note 2 for details)

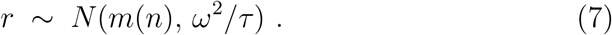

Our goal is now to find a solution of how such a dynamic perceptual system should operate under limited resources. Crucially, we suppose the average value of *m*^2^ is subject to a power constraint, that is to be within some finite bound

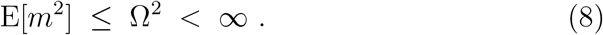

This bound on the amount of variation in the drift limits the precision with which different stimuli can be perceived for any given *τ*. The value of *τ* is assumed to grow linearly with the viewing time, up to some time bound *t*_*max*_,

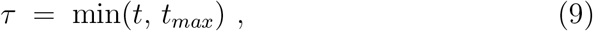

representing a constraint on the amount of time that the decision maker is willing to invest in the accumulation of evidence. The latter bound constrains the degree to which precision can be increased by further increases in viewing time.

The definition of this optimization problem with constraints effectively states that *r* can be seen as the output of a *Gaussian channel* with input *m* [27] that depends on the input stimulus *n*; hence the problem of optimally choosing the function *m*(*n*) is equivalent to an optimal encoding problem for a Gaussian channel (S1 Appendix–Supplementary Note 2).

The capacity *C* of such a channel is a quantitative upper bound on the amount of information that can be transmitted regardless of the encoding rule, which is equal to (S1 Appendix–Supplementary Note 3)

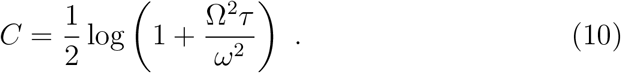

Note that in our model the channel capacity *C* grows as a logarithmic function of *τ* because the correlation of successive increments in the encoding by a Brownian motion prevents the information content from growing linearly in proportion to such increments.

Here we assume that the goal is to design a capacity-limited system that minimizes the mean squared error (MSE) of the estimate 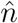 when *n* is drawn from a log-normal prior distribution (i.e., the same objective function stated in the previous section, see Fig. 1). It is possible to show that in our optimization problem, which assumes a channel with “power transmission” constraint Ω^2^, the optimal drift function is

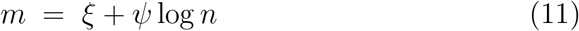

with

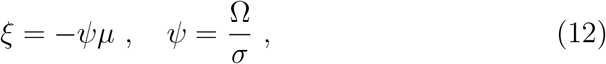

and the encoding noise *ν* in Eq. 2 is given by (see S1 Appendix–Supplementary Note 2 for proof)

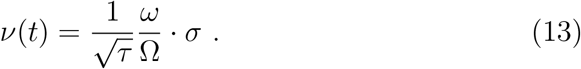

That is, encoding precision grows with viewing time *t* (until reaching the bound *t*_*max*_ if the stimulus is presented long enough). Recall that *σ* is the variance of the log-normal prior, and therefore the solution reveals that the likelihood is independent of parameter *μ* of the log-normal prior distribution, but depends on the second moment of this prior distribution and viewing time *t*. Defining *R* ≡ Ω*/ω*, th<u>e</u> noise of numerosity encoding is given by *ν*(*t*) = 1*/G*, where 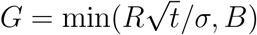and *B* a maximum biologically allowed bound on sensory precision related to 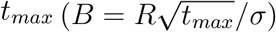. Note that the two information bounds affect the model differently. The higher is Ω, the more information gathered per time unit, whereas *B* captures the maximum amount of information that can be gathered. Notice that this solution implies a multiplicative trade-off between Ω, *ω*, and *t* (similar to the standard DDM). However, this relation may look different under other assumptions (e.g., non-uniform noise in the encoding space).

**Fig 1.**
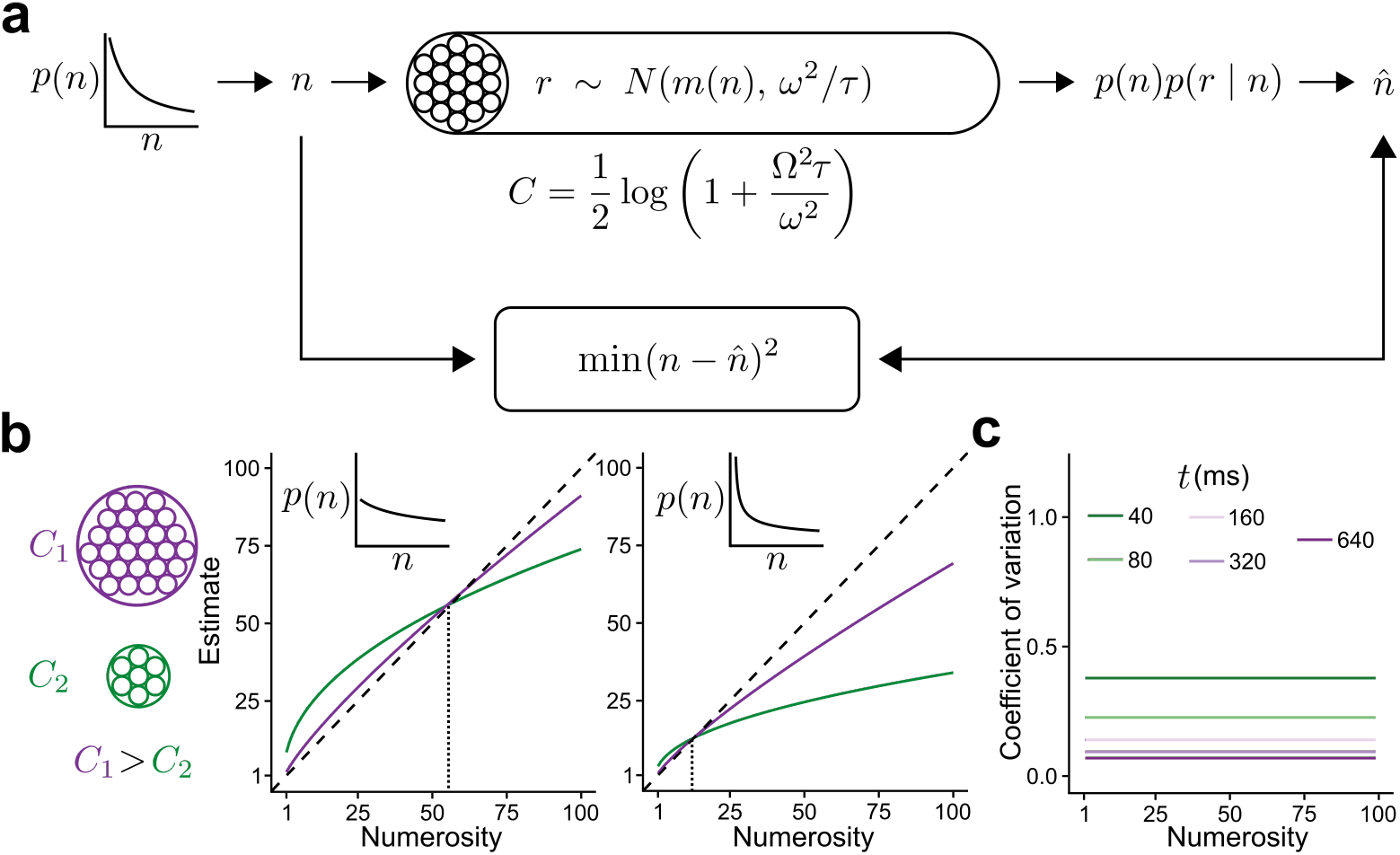
Overview of the SEB model. **a)** Schematic description of the SEB model. A numerosity *n* is drawn from a prior distribution *p*(*n*). The observer has a limited capacity *C* to represent the numerosity. The internal representation *r* is a random draw from a Gaussian distribution, the mean of which depends on *n* but the variance does not. The observer then infers the estimate 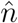 based on the representation *r* and the prior distribution of *n* as to minimize the MSE between the estimate and the numerosity. **b)** Illustration of the predictions for an observer with a high (purple) or low (green) channel capacity where *n* is drawn from a distribution with a high (left) or low (right) variance. All curves exhibit overestimation for lower numerosities and underestimation for higher numerosities. However, these biases are reduced in the case of high capacity. The crossover point between under- and overestimation increases with the variance of the numerosity distribution. **c)** Illustration of the coefficient of variation (i.e., 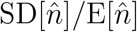) for different capacities. The coefficient of variation is independent of the numerosity and decreases with capacity, which is dependent on the viewing time *t*.

These results lead to the following predictions from our model: (i) 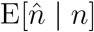 is a concave function of *n* with overestimation for small numbers (when these are not so small that the discreteness of available responses leads to nearly-deterministic responses), but underestimation for large numbers (Fig. 1b and S1 Appendix–Supplementary Note 2). (ii) The crossover point from overestimation to underestimation changes as a function of the numerosity range and prior variance (see S1 Appendix–Supplementary Note 4). In addition, the concavity of 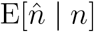 depends on the amount of resources available to perform the numerosity estimation task. This prediction was clearly confirmed in a previous empirical work [4]. (iii) Because of the discreteness in the set of responses, there is predicted to be little variability in responses in the case of small enough numbers. This may look in principle as a subitizing-like behavior for small numbers. However, SEB does not predict subitizing in principle. Subitizing-like behavior in SEB results from smaller estimation biases and variability by the observer which may be experimentally imperceptible after rounding to generate a discretized response. (iv) For numbers beyond the subitizing-like range, based on the properties of the log-normal distribution, it can be shown that the coefficient of variation (S1 Appendix–Supplementary Note 1)

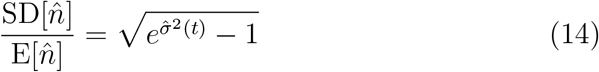

does not depend on the input numerosity *n*, thus delivering the property of scalar variability, irrespective of *n* [5], but here we show that this coefficient will depend on time exposure *t*, with the predicted constant co<u>efficient</u> of variation decreasing as *t* gets larger proportionally with 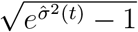 (Fig. 1c).

### A thermodynamically inspired model of bounded rationality

Here we briefly introduce a popular approach to studying systems with bounded capacity across domains in human cognition and machine learning: a thermodynamically inspired formalization where information processing is modeled as changes from a default state, which come at some energetic cost, that can be quantified by differences in free energy. This class of models can be applied for the case where an observer intends to minimize some form of expected loss (here we study the case of estimation error minimization), subject to information constraints [17]. More formally, let 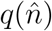 be a default state (distribution) over possible responses 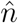 in a given environment or context. When presented with a stimulus *n*, the resource-constrained observer attempts to transform the initial state *q* into a new state of possible responses 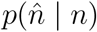. This transformation of states can be modeled as the optimization of the free energy functional

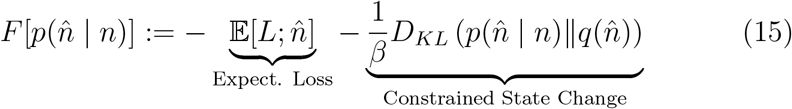

where *L* is a loss function, for instance, the squared error 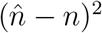. The second term is the Kullback-Leibler divergence between *q* and 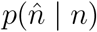, where *β* trades off the relative importance of changing from the default state *q*, thus determining the resources that the observer invests in the estimation task. The goal is to find the optimal distribution of responses

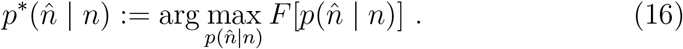

The optimal distribution of responses in this variational problem has an analytical solution of the form

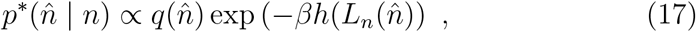

where *h* is a function of *L* and potentially other elements incorporated in the expected loss function in Eq. 15.

### TIM applied to numerosity estimation

A recent work applied a model from the TIM family to study a resource-constrained model of human numerosity estimation [21]. This is also a formulation of how the distribution of reported numerosity estimates 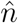 of a stimulus magnitude should vary depending on the true stimulus *n*. This can be stated generally as the hypothesis that conditional on *n* the response distribution 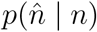 is the probability distribution over a set of possible responses *N* that minimizes the mean squared error (MSE), subject to the constraint

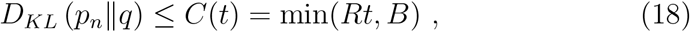

where *C*(*t*) is a positive bound that depends on the amount of time *t* for which the stimulus is presented. This formulation can be interpreted as a model in which errors in the observer’s responses can be attributed to a “cost of control” of the responses: it is difficult for the observer to give responses different from the default state *q*, though their response distribution to the individual stimulus *n* is optimal given a constraint on the possible precision of their responses.

Similar to our SEB model, in TIM it is assumed that perception extracts information linearly in time at a rate *R* until an overall capacity bound *B* is reached. The goal is to find the distribution of numerosity estimates 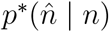 that minimizes the mean squared error

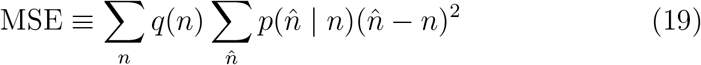

under the constraint given in Eq. 18.

The optimization problem described above yields the following analytical solution [21]

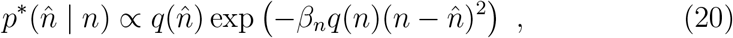

where *β*_*n*_ is chosen to satisfy the bound in Eq. 18. Note that this solution has the familiar form obtained in Eq. 17 with *L* as the loss function 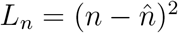.

While this solution is usually linked to a bounded-rational Bayesian computation (given the observation that the default distribution *q* is multiplied by a function of *n* given 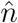), here we clarify that this solution does not correspond to a Bayesian inference process with noisy sensory percepts. In fact, the TIM formulation assumes that the perception of the sensory stimulus *n* is noiseless, and all the variability observed during the estimation process is related to the cost of acting accurately, that is, a cost in the precision of response selection when shifting away from the default state. Note that this is fundamentally different from the SEB model, in which all the estimation variability is attributed to noisy sensory encoding.

### Overview of the constraint parameters of the SEB and TIM models

One of our goals is to formalize and make transparent the different elements that play a role in a noisy information transmission process under our model specification, namely, (i) time, (ii) precision of instantaneous information processing, and (iii) energy required to transmit decodable information.

On the one hand, the constraint *B* is similar in spirit to the DDMs applied to cognitive processes, where the decision maker is willing to invest a maximum amount of time in processing information due to opportunity costs, in principle irrespective of how precise the instantaneous information processing is. Thus, formally, the time invested in acquiring information is given by

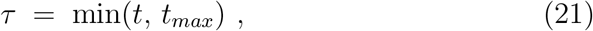

with *t*_*max*_ defining the maximum information bound 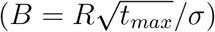.

On the other hand, the parameter Ω in our work imposes a constraint on the cost of a decodable transmitted message considering deviations from a status quo state, where energy needs to be injected to transmit decodable information (which in our model specification is given by *m*(*n*), for any given *τ*. However, the message *m* is not noiseless, where instantaneous processing noise is given by *ω* (otherwise one could make the length of *m* infinitesimally small). This means that there exists a natural trade-off between the fidelity of information processing, how much energy the decision maker is willing to (or can) “pay” to disentangle information, and also how much time should be invested in this process. In addition to this, there is an objective that we assume the decision-maker would like to optimize for: minimize the mean squared error. Here it is important to emphasize two points: (i) specifying the model in this way makes transparent the different limitations that the decision-maker must trade-off; (ii) the constraint *B* is not a necessary requirement to obtain the optimal solution. It is set following the common knowledge that processing time leads to opportunity costs, in principle irrespective of information processing precision. These constraints will almost surely trade-off in the presence of imprecise information processing (as is also the case in n-alternative choice DDMs). Should time be allowed to be infinite then decoding would be nearly perfect even with imprecise instantaneous information processing (Equations 7 and 10); analogously in n-alternative DDMs with noisy drifts, bounds would be infinitely large, thus converging to errorless decisions.

These constraint assumptions are also present in the TIM:

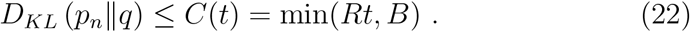

Here the constraint is similarly given by two parameters: a linear information “processing” rate *R*, and (ii) a bounded rate of information processing *B*. However, as we discuss in detail in our work, the TIM specification does not consider noisy encoding.

### General similarities and differences between SEB and TIM

We elaborated an illustrative example that allows the predictions of the two models to be solved analytically, thus allowing us to understand the key differences between them (S1 Appendix–Supplementary Note 5). These analyses reveal some similarities between the predictions of the two models, however, there are also notable differences. First, while both models predict that biases decrease in general for larger viewing times, this decrease occurs sooner for SEB than TIM. Second, for a given input stimulus *n*, the two models do not imply that 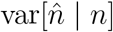 co-varies with the bias in the same way. As the viewing time *t* goes to zero, the Bayesian model implies that the variance should fall to zero; TIM implies that this is the case in which estimates should have the highest variance (equal to the variance of the prior distribution).

These analytical insights were studied over all possible responses in the continuous space and do not directly apply to numerosity estimation in the discrete space (S1 Appendix–Supplementary Note 5). Therefore, we conducted numerical analyses to study whether the same signatures emerge in SEB and TIM when the solutions are restricted over the space of positive integers and to give some intuition by visualizing the differences.

As expected, both models predict that biases decrease in general for larger viewing times, and mirroring the results of the analytical solution, this decrease occurs sooner for SEB than TIM (Fig. 2, left panels).

**Fig 2.**
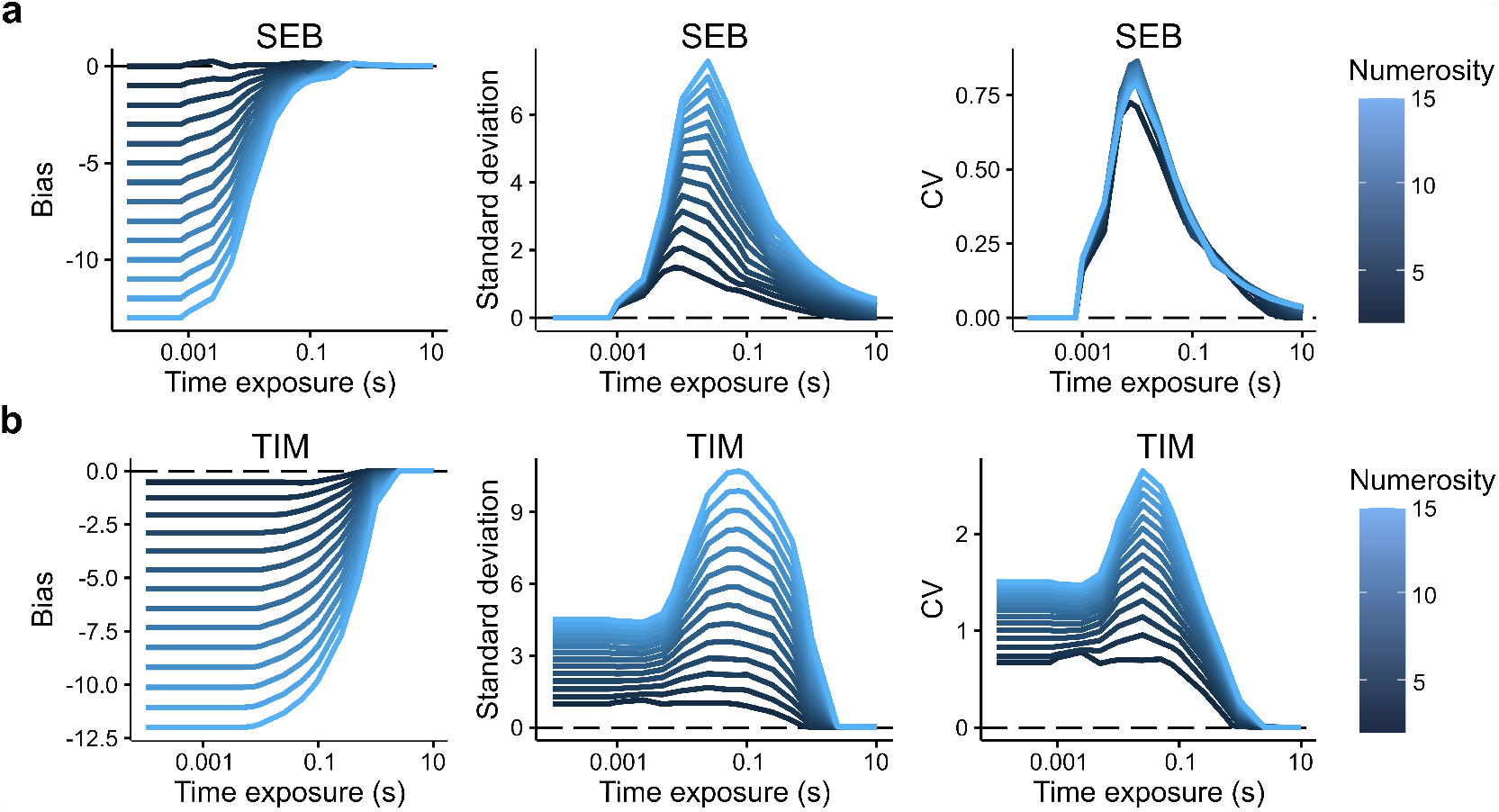
General similarities and differences between SEB and TIM. **a)** Computation of the bias (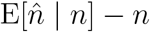, left panel), standard deviation (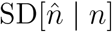, middle panel), and the coefficient of variation (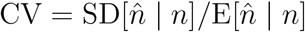, right panel) as a function of different time exposures *t* for different numerosities *n* (color scale of the solid lines) in the SEB model. Although we proved that the CV in SEB does not depend on numerosity (14), notice that in the simulations the CV does vary slightly. These slight differences are the result of rounding the expected value of the posterior to the closest integer for the discrete SEB model. For smaller integers, the CV will become more affected by these rounding errors, i.e., due to a slight overestimation of the Bayesian decoder for smaller numbers, the expectation will slightly increase thus dominating the CV ratio.

However, a fundamental difference between the two models is that as *t* goes to zero, the SEB predictions SEB fall to zero, but this is not the case in TIM where the predicted variability of estimates is clearly larger (Fig. 2, middle panels).

Finally, the computation of the coefficient of variation 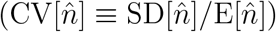 reveals that in SEB this metric is nearly identical for all numerosities *n* irrespective of time exposure *t*, thus reflecting the scalar variability effect (Fig. 2a, right panel). In TIM, however, the scalar variability phenomenon is absent irrespective of time exposure *t*. These differences make the two models different and identifiable and generate somewhat different qualitative predictions.

### Efficient numerosity estimation under limited time

We now compare TIM with SEB models using the experimental data of a pre-registered study provided in previous work [21] (see Methods). In brief, on each trial, between 1 and 15 dots were flashed, followed by a noise mask. The participants were then asked to type their estimation of how many dots were displayed. There were three between-participant experiments (n=100 per experiment) that manipulated available stimulus information (variable exposure time: *t* ∈ [40, 80, 160, 320, 640] ms) and different ways of controlling non-numerical properties of the stimuli: the average dot size (experiment 1), surface density (experiment 2) or surface area (experiment 3) of the dots.

To fully constrain inference solely to the normative solutions of stimulus exposure derived above for both SEB and TIM, we fixed the prior distribution before fitting the behavioral data to a prior equivalent of the form 1*/n*^*α*^ power-law. It has previously been argued that the prior probability of how often numerosities are encountered and represented roughly follows a 1*/n*^*α*=2^ power-law distribution [24, 25]. Thus, a priori, we choose *α* = 2, following the same assumption adopted in previous work [21]. By fixing such ecologically valid prior, we alleviate the critique of allowing an arbitrary choice of prior and likelihood functions to fit inference models to the data, as a consequence of which it is sometimes argued that their predictions are potentially vacuous [28]. Nevertheless, it is well possible that each individual has learned their own distribution during their lifespan [29, 30]. Therefore, we also considered a more flexible class of models where we allowed the parameters of the prior distribution to be free parameters alongside the capacity constraint and capacity bound.

We considered two possible ways of inferring the numerosity estimates based on the SEB approach (methods): (i) using the analytical solutions over the continuous positive real line, and (ii) using discrete encoding and decoding restricted to the positive integer numbers, thus similar in nature to the TIM specification. Finally, we considered a guessing rate *g* in the model fits, which assumes that on *g* proportion of trials, participants were distracted and had no information about the number of dots in the display, meaning that their estimate was effectively a random sample from their prior. Thus, both numerosity estimation models SEB and TIM have exactly the same degrees of freedom (the capacity constraint, capacity bound, and *g*), in addition to the prior parameters in the flexible class of models.

### Quantitative model comparison

For each experiment where stimulus presentation time *t* was manipulated, we fit both types of model to the data of each participant (Methods). In parameter recovery exercises we found that all model parameters are identifiable and this is also confirmed by the weak relationship between parameters across participants (Supplementary Fig. 1). We first examined the restricted models where the prior is fixed 1*/n*^2^. For experiment 1, we found that the difference in Akaike information criterion (AIC) favoured SEB, where the continuous version of SEB had a clear advantage over TIM: ∆AIC=1472 [95%-CI 570-2553] in favor of SEB (paired t-test: *T* (99) = 2.86, *p <* 0.01, *d* = 0.29. For experiment 2 (dot density controlled), the difference in AIC is 5284 [95%-CI 4185-6690] in favor of SEB (*T* (99) = 8.70, *p <* 0.001, *d* = 0.87). For experiment 3 (dot area controlled), the difference in AIC is 2316 [95%-CI 1218-3686] in favor of SEB (*T* (99) = 3.61, *p <* 0.001, *d* = 0.36, see Fig. 3a). In addition, the SEB continuous model provided better fits than its discrete version (*T* (99) ≥ 8.64, *p <* 0.001, *d >* 0.86 ∆ AIC ≥997).

**Fig 3.**
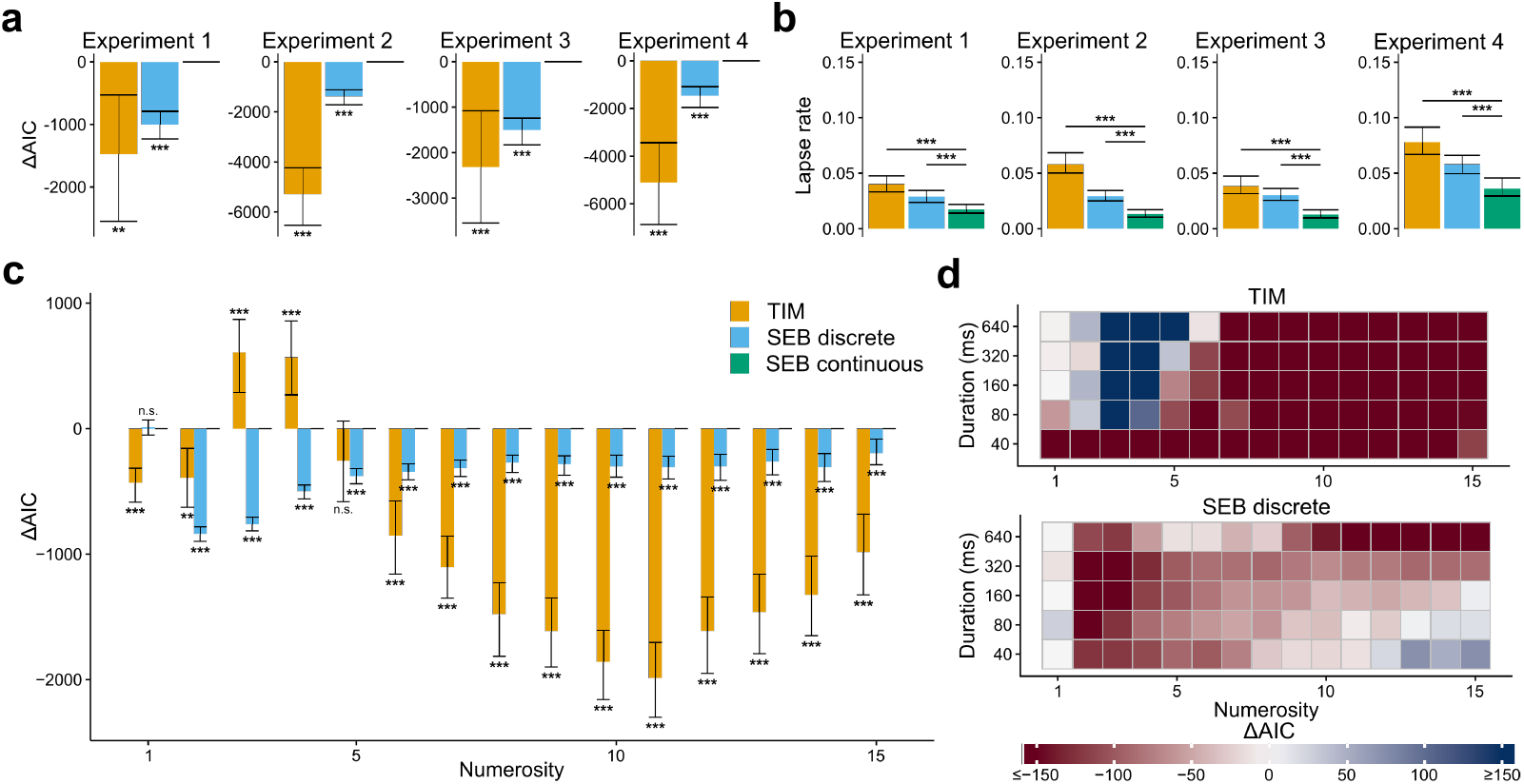
The SEB model quantitatively outperforms the TIM model when the prior parameters are fixed. **a)** Difference in AIC between the SEB continuous model (green) and the TIM model (orange) or the SEB discrete (blue). The ∆AICs were computed for each participant and summed. The error bars represent the 95% confidence interval based on bootstrapping of the participants’ ∆AICs. The SEB continuous model outperforms both the TIM model and the SEB discrete model. **b)** Average guessing rate parameter per participant for each model and experiment. The error bars represent the 95% confidence interval based on bootstrapping of the participants’ guessing rate. The guessing rate of the SEB continuous model is lower than the TIM model and the SEB discrete model. These results indicate that less variability is associated to lapses of attention in the SEB continuous model, which suggests a better fit to behavior. **c)** Difference in AIC between the SEB continuous model and the TIM model or the SEB discrete model for each numerosity. The error bars represent the 95% confidence interval based on bootstrapping of the participants’ AICs. The SEB continuous model outperforms the TIM model except for numerosities 3, 4 and 5 and the SEB discrete model for all numerosities except numerosity 1. **d)** AIC differences between the SEB continuous model and the TIM model (top) and the SEB discrete model (bottom) for all experiments shown for different numerosities and levels of sensory evidence (stimulus presentation duration or contrast). Duration values are assigned to Weber contrasts of experiment 4 for pooling purposes (40ms–10%, 80ms–20%, 160ms–40%, 320ms–80%, 640ms–160%). The SEB continuous model outperforms the TIM and SEB discrete models for most numerosities and levels of sensory evidence.

Previous theoretical and empirical work suggests that two ways in which the amount of information available to process information can be studied are by manipulating time exposure and also by changing stimulus contrast [13]. Thus, we also considered this alternative way of manipulating sensory reliability, which should affect the channel capacity transmission (see Eq. 10). To test this, we analyzed data of a numerosity estimation experiment, where in each trial the visual contrast of numerosity was manipulated at a constant presentation time (n=100 participants, experiment 4, Methods). We found that also in this experiment the SEB-continuous model fits the data better than TIM (∆*AIC* = 5106; [95%-CI 3452-6880] (*T* (99) = 5.79, *p <* 0.001), *d* = 0.58, Fig. 3a) and the discrete version of SEB (∆*AIC* = 1453; [95%-CI 1059-1907] (*T* (99) = 6.69, *p <* 0.001, *d* = 0.67)).

To make sure that the overall quantitative differences were not driven by a few numerosities, we computed the difference in AIC for each numerosity and each model. We found a significant interaction models*numerosity of the ∆AICs (*F* (28, 16758) = 7.84, *p <* 0.001) with posthoc tests revealing that this effect was more pronounced for higher numerosities (SEB continuous vs. TIM: paired t-tests *p <* 0.001 for numerosities *n >* 5, Fig. 3b) and also for *n* ∈ [1, 2] (paired t-tests *p <* 0.01). The relative advantage of the TIM model for *n* ∈ [3, 4]) at large presentation times *t* might be explained by the fact that smaller numerosities are close to the subitizing range and therefore most of the posterior density mass is concentrated around the input *n*, which is better explained by the TIM model as this model has a tendency to subitize more strongly at small numerosities [21]. Interestingly, for *n* ∈ [1, 2], the Bayesian model predicts noisier estimations (in particular for smaller exposure times *t*) which are not supported by the TIM, with the AICs favoring the former.

Additionally, we inspected the AIC differences split by both numerosity and sensory evidence (time or contrast), finding a similar pattern, but the differences were larger for small levels of sensory evidence. Thus, SEB appears to be more sensitive to capturing behavior for stimuli generating higher noise levels in the encoding operations.

Moreover, we compared the guessing rates *g* between the two kinds of models. Guessing rates can capture unassigned variance in misspecified models, thus we conjectured that a relatively smaller value of *g* would provide further evidence for better mechanistic fits captured by the best model. While the guessing rates are overall small (suggesting that the amount of distractions during task performance was minimal), we found that guessing rates were systematically smaller in the SEB model (*T* (99) *≥* 5.75, *p <* 0.001, *d >* 0.58 for each experiment, Fig. 3b and Supplementary Table 1). Thus, while the effects of distraction are estimated to be relatively small in both models, our analyses provide a clear indication that potentially unassigned variance due to distraction is lower in the SEB model relative to TIM. We also note that the information bound parameter *B* can be fit for both models which is empirical evidence that this information bound is present in this experimental setup, even for relatively short presentation times (640ms).

We repeated the same set of analyses treating parameters of the log-normal prior as free parameters. The results of these analyses mirrored the initial analyses. That is, (i) we found that the SEB model fit the data better than TIM in all four experiments (*T* (99) *≥* 3.54, *p <* 0.001), Fig. 4a), (ii) the continuous version of SEB performed better in general than its discretized version (Fig. 4a) and (iii) the guessing rates were significantly smaller in the SEB model than the TIM model for experiments 2 and 3 (*T* (99) *≥* 2.95, *p <* 0.01 but not for experiments 1 and 4, Fig. 4b).

**Fig 4.**
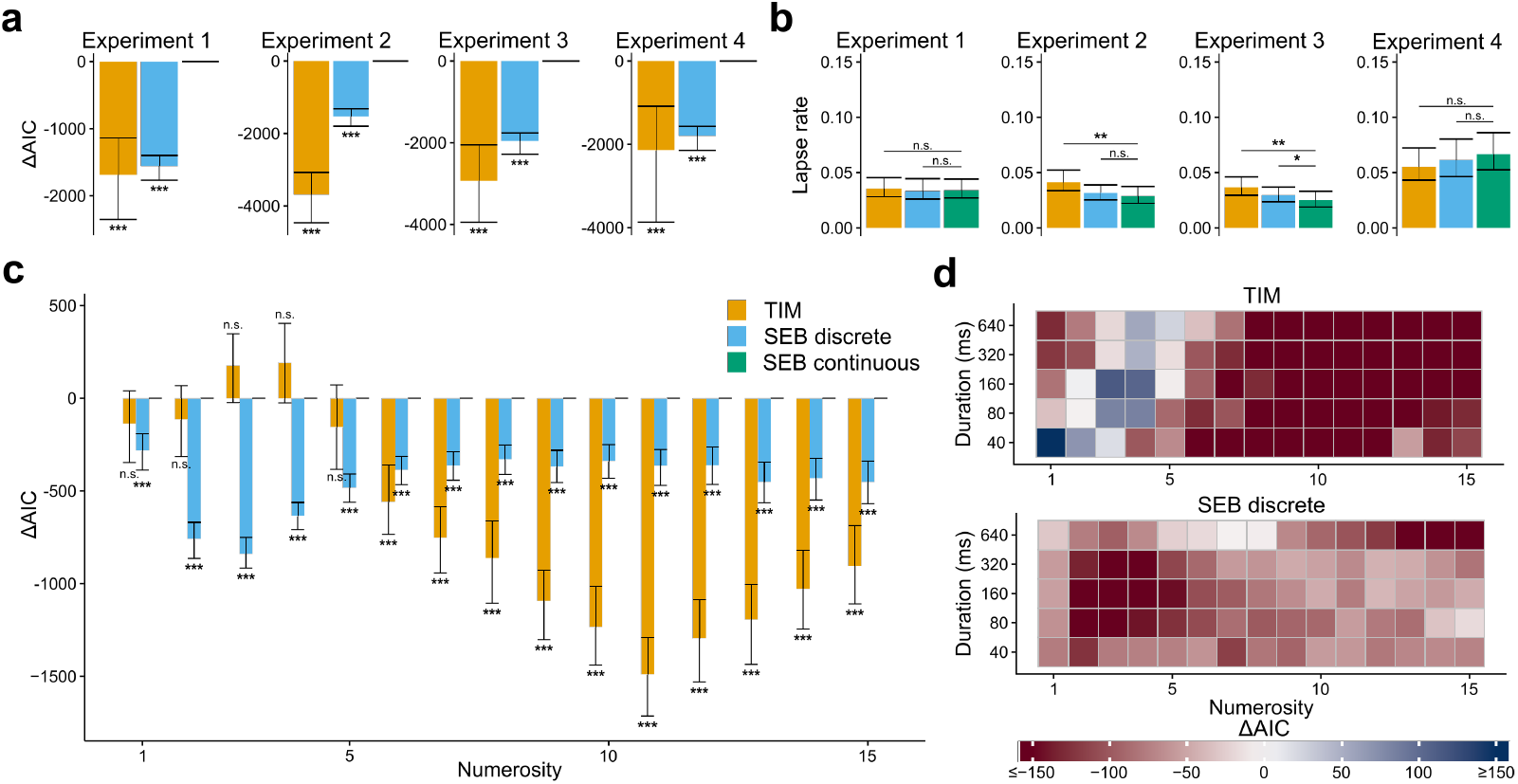
The SEB model quantitatively outperforms the TIM model when the prior parameters are free. **a)** Difference in AIC between the SEB continuous model (green) and the TIM model (orange) or the SEB discrete (blue). The ∆AICs were computed for each participant and summed. Error bars represent the 95% confidence interval based on bootstrapping of the participants’ ∆AICs. The SEB continuous model outperforms both the TIM model and the SEB discrete model. **b)** Average guessing rate parameter per participant for each model and experiment. Error bars represent the 95% confidence interval based on bootstrapping of the participants’ guessing rate. The guessing rate of the SEB continuous model is lower than the TIM model for experiments 2 and 3 but not for experiments 1 and 4 and the SEB discrete model for experiment 3 but not for the other experiments. **c)** Difference in AIC between the SEB continuous model and the TIM model or the SEB discrete model for each numerosity. Error bars represent the 95% confidence interval based on bootstrapping of the participants’ AICs. The SEB continuous model outperforms the TIM model except for numerosities 1 to 5 and the SEB discrete model for all numerosities. **d)** AIC differences between the SEB continuous model and the TIM model (top) and the SEB discrete model (bottom) for all experiments shown for different numerosities and levels of sensory evidence (stimulus presentation duration or contrast). Duration values are assigned to Weber contrasts of experiment 4 for pooling purposes (40ms–10%, 80ms–20%, 160ms–40%, 320ms–80%, 640ms–160%). The SEB continuous model outperforms the TIM and SEB discrete models for most numerosities and levels of sensory evidence.

The next question to ask is whether the models with free prior parameters outperformed the models with the prior fixed to 1*/n*^2^. We found that for each model considered here, the models with free prior parameters outperformed their corresponding version with fixed parameters (*T* (99) *≥* 5.62, *p <* 0.001, Supplementary Table 2). Additionally, to account for population variability in the quantitative metrics between participants across all models considered here, we applied a Bayesian Model Selection which revealed that the Bayesian model with free prior parameters is clearly favored relative to all the other models for experiments 1, 2 and 3 (*P*_xp_ *>* 0.99 for each experiment) but equally favored to the TIM model with free prior parameters for experiment 4 (*P*_xp_ = 0.50). These results allow us to conclude two important points. First, variability in the prior parameters of the prior distribution is key to more accurately explaining human numerosity estimations. Second, our results provide a clear indication that the effects of temporal time exposure are better captured by the noisy encoding model (SEB) relative to an action control-like model (TIM).

### Qualitative predictions

We first examined the qualitative features of scalar variability in both data and the predictions of the SEB continuous and the TIM models with free prior parameters. For each numerosity value, we computed the coefficient of variation 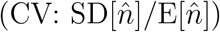. We found that the empirical data follows the previously observed properties of scalar variability for numerosities greater than 4 (i.e., a flat CV irrespective of numerosity and sensory evidence), with a slight systematic increase of CV for smaller numbers (Fig. 5a left). This relative CV increase for small numbers could be explained by the presence of small guessing rates *g* which have a greater impact on the CV for small *n*. We found that the SEB model accounts for these qualitative observations (Fig. 5a middle), however, the TIM model generates slightly different predictions (Fig. 5a right).

**Fig 5.**
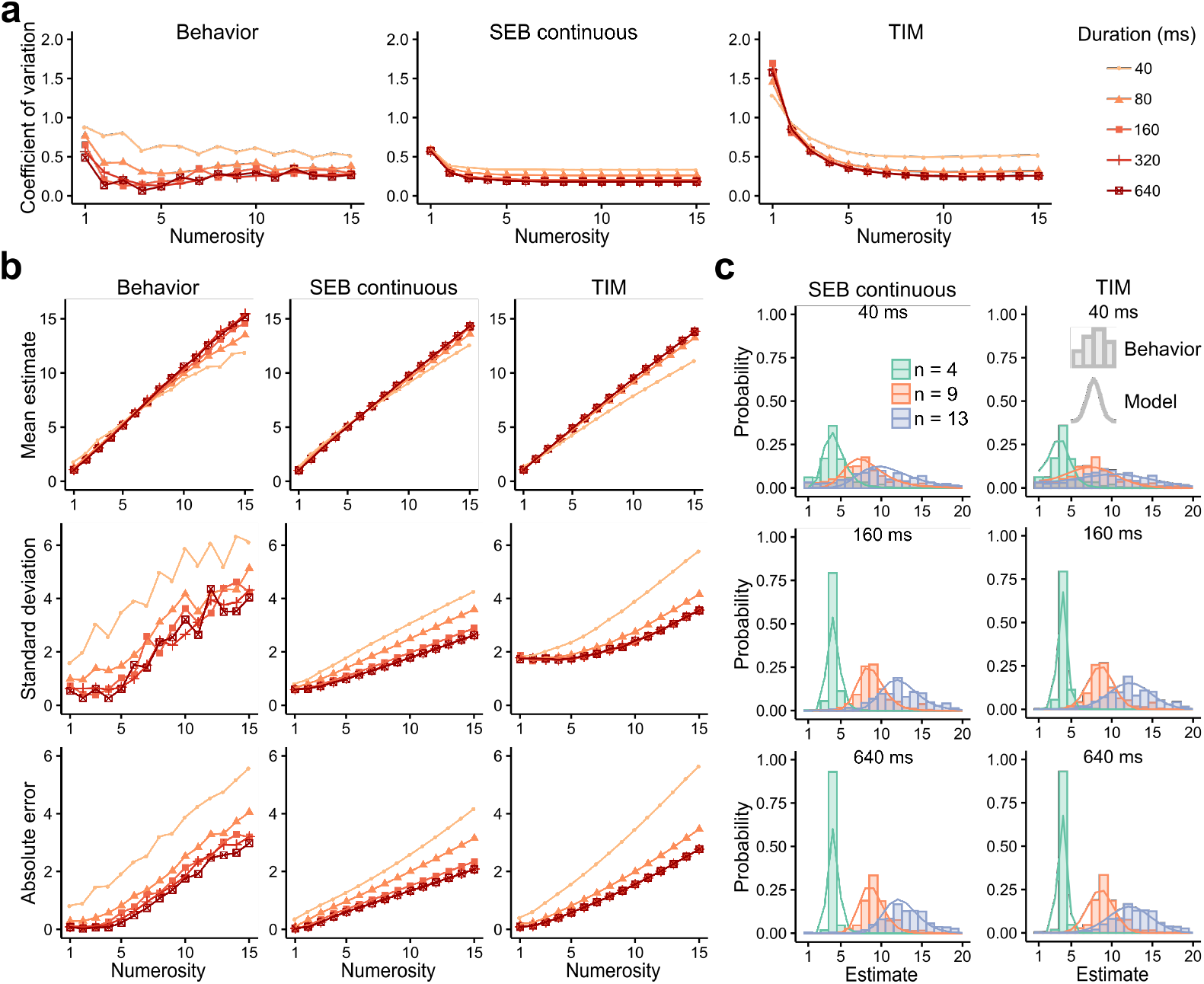
The SEB continuous model with free prior parameters qualitatively explains behavior. **a)** Coefficient of variation 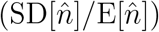 of the behavior data (left) and predicitons of the SEB model (middle) and TIM model (right) using a prior with free parameters for different numerosities and stimulus presentation duration. Predictions were performed by taking for each parameter the value with this highest density across participants. Duration values are assigned to Weber contrasts of experiment 4 for pooling purposes (40ms–10%, 80ms–20%, 160ms–40%, 320ms–80%, 640ms–160%). The TIM model predicts a higher CV for lower numerosities. This feature is not present in the behavior data nor the SEB predictions. **b)** Mean estimate (top), standard deviation (middle) and absolute error (bottom) of the behavior data (left) and predictions of the SEB model (middle) and TIM model (right). **c)** Posterior distribution of estimates to numerosities 4 (green), 9 (red) and 13 (blue) of the SEB (left) and the TIM (right) model for different stimulation presentation durations (40ms (top), 160ms (middle), 640ms (bottom)). Behavior of participants is shown as histograms.

We found that patterns of estimation biases and variability during numerosity estimation as a function of sensory evidence were in general more closely captured by the SEB relative to the TIM model (Fig. 5b top and middle panels). As predicted by our analytical analyses (S1 Appendix–Supplementary Note 5) the rate of increase in noise as a function of *n* is larger for the TIM model relative to the SEB model, with the empirical data more closely agreeing with the SEB model. Additionally, given that the TIM model generally requires larger values of guessing rates *g* to explain variance, for small *n* it predicts larger SDs relative to SEB and empirical data (with a similar pattern for the case of the CV, (Fig. 5a). However, there is an exception at 40ms, where TIM captures better the range of the standard deviation (from around 2 to 6). Another point where the TIM model appears to do a better job relative to the SEB model is for the absolute error estimations (Fig. 5b bottom). Subitizing is more pronounced for low numbers in general, and this reduces both biases and errors for *n <* 5. However, beyond the subitizing range and for levels of noise that challenge sensory perception, the SEB model does a better job at capturing all descriptive statistics. To visualize the nature of these differences, the posterior distribution of estimates for both models are shown in Fig. 5c for different numerosities and presentation times.

## Discussion

Our theoretical and empirical tests provide clear evidence that a model of Bayesian decoding of noisy internal representations—which provides a normative explanation for the property of scalar variability and can be parsimoniously connected to a theory of limited informational capacity—provides a better account of numerosity estimation data in humans relative to the alternative TIM model considered here. We emphasize that both models: (i) are optimized for the same assumed objective (minimizing the MSE of the estimates), (ii) can be compared under the same assumption about the prior distribution, and (iii) have identical degrees of freedom. Thus, qualitative and quantitative differences between the two information-theoretical models cannot be explained by differences in model complexity, but instead reflect differences in the mechanistic assumptions of the numerosity processing operations. In particular, it is important to note that assumptions about potential encoding and decoding operations are explicitly stated in the Bayesian model. In contrast, these remain “hidden” in the alternative TIM model.

One of our main goals in the development of our modeling framework was to develop an encoding-decoding model incorporating various aspects of human cognition with many antecedents in the literature, which include Brownian motion during evidence processing over time [15] and logarithmic internal representation of numerical quantities [16]. While our proposed model accounts for key qualitative features of the human behavioral data with minimal degrees of freedom, we do not claim that the log-encoding model necessarily accounts for all aspects of numerosity estimation behavior. Indeed, the encoding and decoding strategies that humans and other animals use need not be the same in all contexts [31]. It is equally possible that numerosity processing mechanisms depend on the task at hand, and draw upon an ensemble of strategies that optimize performance under different situations [32, 33]. For instance, in future work, it will be interesting to investigate whether situations that involve explicit numerosity estimation vs. discrimination rely on similar or distinct encoding strategies and inference processes.

We assumed that participants employ a log-normal (or power-law) prior, however, it is important to note that the numerosities presented to the participants were drawn from a uniform distribution. We thus implicitly assumed that participants did not rapidly adapt their encoding operations, which might be a reasonable assumption given that participants were not exposed to the new prior for an extended period of time. However, in one version of the model fits we allowed the parameters of the prior to be adjusted, resulting in non-uniform distributions, which at the very least suggests that participants did not adapt to a uniform prior.

A natural consequence of our theory is that the SEB model parsimoniously endogenizes parameters of the prior distribution in its encoding operations. A testable prediction is that larger prior distribution ranges should lead to more noisy estimates and therefore poorer discriminability for a given capacity bound. This prediction is confirmed by a recent study where it is shown that human participants adapt their numerosity sensitivity for different numerosity ranges, with important implications for risk behavior [34]. Thus two of the key predictions of our theory hold: for a fixed capacity bound sensory reliability should change as a function of (i) time exposure to the sensory stimulus as shown in this study, and (ii) the range of the prior distribution [34].

Additionally, our model predicts that the crossover point from overestimation to underestimation should change as a function of the numerosity range. In this work, we only present data with a fixed range of 1 to 15, thus we cannot test this prediction. However, a previous study using larger numerosity ranges (e.g., up to 30 or 100) found that the cross-over point is larger for wider numerosity ranges, and crucially, the degree of over- and under-estimation depended on the attentional resources dedicated to numerosity estimation [4]. This result is again in line with the qualitative predictions of our model. Future research could test these predictions quantitatively.

The general working framework of SEB is a Bayesian decoder where the likelihood function optimally endogenizes information in the prior distribution for a given capacity constraint and stimulus exposure time. Here it is important to emphasize that this framework is not restricted to a specific form of the prior. Also, the formulation of processing time, the information processing constraint, and the objective are specified in a general form. However, a fundamental aspect of the SEB model is the specific form of the drift-diffusion term which employs an affine (log-)linear transformation of the input stimulus. The question here is whether this affine function is valid and generalizes for the case of numerosity estimation when it is assumed that the prior distribution changes (i.e., ceases to be power-law or log-normal). We argue that unless there is extensive training over long periods (e.g., many days of ecologically valid adaptation in the absence of any power-law or log-normal distributions) the encoding function may remain of the family of an affine (log-)linear of power law transformation. While we acknowledge that this argument needs to be tested empirically, recent work appears to support this notion: Prat-Carrabin and Gershman [35] show that when either the prior or objective functions (or incentives) are manipulated during a numerosity estimation task, and even if behavior shows signatures of such adaptation, modeling analyses suggest that subjects’ responses feature in all cases logarithmic encoding while the Bayesian decoder takes into account the prior distribution and objective function (incentives). Nevertheless, this form of the encoder might not be optimal and may need to be adapted to the prior distribution and objectives of the organism to achieve efficiency. This idea was studied in previous work both theoretically and empirically where it is shown that if organisms can adapt their encoding functions, they must do so at the earliest stages of sensory processing [36, 37], otherwise, information that is lost in the early sensory processing streams cannot be recovered via downstream operations [37, 38].

In addition, given the relatively short stimulus presentation times, we assumed that the participants gather information about the stimulus until they reach the time bound *t*_*max*_ or the stimulus disappears. Other specifications of perceptual decision-making problems include endogenous stopping, in which case the observers decide by themselves when to stop gathering information and respond. Although this question is out of the scope of this study, future researchers could build on our model and specify a utility function which takes into account both the rewards for accurate answers and costs related to the time of the decision.

Our model is agnostic about the biological meaning of its parameters. Future research could try to relate them to neural processes. We speculate that the bound on the drift rate Ω could be related to the information capacity of sensory areas (for example how many neurons are used to represent a stimuli or how much precision can these neurons use). This contrasts with the information bound *B*, related to the maximum amount of information that can be represented, which could be related to neurons resources in higher cortical areas such as the dlPFC as well as premotor areas.

Taken together, our findings suggest the fruitfulness of studying optimal models with resource limitations, which can serve as a departing point to understand the neuro-computational mechanisms underlying human behaviour without ignoring the fact that biological systems are limited in their capacity to process information [36, 37, 39, 40]. This highlights that understanding behavior in terms of its objectives while taking into account cognitive limitations, alongside encoding, decoding, and inference processes is likely to be essential to elucidate the mechanisms underlying human cognition.

## Materials and methods

### Participants, data, and experiments

In this work we re-analyzed the data of experiments collected in previous work [21]. In brief, on each trial, between 1 and 15 dots were flashed, followed by a noise mask. The participants were then asked to type their guess of how many dots were displayed. The participants were recruited and carried out the experiment online. There were three between-participant experiments (n=110 per experiment) that manipulated available stimulus information (variable exposure time: *t* ∈ [40, 80, 160, 320, 640] ms) and different ways of controlling non-numerical properties of the stimuli: the average dot size (experiment 1), surface density (experiment 2) or surface area (experiment 3) of the dots.

We also studied a fourth experiment (n=110) in which time exposure *t* was fixed across trials, but instead display contrast of the dot arrays was varied from trial to trial (experiment 4). In this experiment, the colors of the dots varied between the background (grey) and pitch black, by Weber contrasts of 10%, 20%, 40%, 80% and 160%, at a constant presentation time of *t* = 200 ms.

Each participant was presented with each combination of numerosity and sensory evidence twice for a total of 150 trials per participant.

### Models

Here we fit the two families of models described in the main text to the data of each participant: (i) We fit the SEB model assuming a log-normal prior with power parameter *α* = 2. We fit a continuous version of the model based on the analytical solutions derived in S1 Appendix–Supplementary Notes 1-2, and a discrete version of this model based on numerical simulations. (ii) Following the procedures of previous work [21], we fit the TIM model assuming a power-law prior with power parameter *α* = 2. For both families of models, we also fit a version were the parameters of the log-normal prior were allowed to be free parameters. We also note that analytical solutions in SEB were derived in the continuous space due to mathematical tractability (S1 Appendix–Supplementary Notes 1-3). Thus, in order to define the likelihood function of this model in the integer space, we normalized the log probability of estimators (Eq. 31) in the integer range *n* ∈ [1, 2, 3, …, 100]. Note that both SEB and TIM have exactly the same degrees of freedom (*R, B*, and *g*), where *g* is a guessing rate based on the probability of randomly drawing a value from the default distribution.

### Quantitative and qualitative analyses

Participants who completed less than 90% of the trials were excluded. Similar to previous work [21] we selected the 100 best participants for each experiment. In addition, trials in which the participant’s response was 10 times higher than the presented numerosity or the response time was superior to 10s were excluded. This additional data cleaning leads to the rejection of 142 trials out of 14,997 for experiment 1, 143 out of 14,993 for experiment 2, 172 out of 15,000 for experiment 3 and 187 out of 15,000 for experiment 4. Each model was fit individually to each participant using the DEoptim package [41] in the statistical language R [42] with a number of iterations set to 100. The limits for the parameter search space were set to (0.1,200) for *R*, (0.1, 20) for *B* and (0.0001, 0.5) for *g*. In the models where the prior was free, the search space of the prior parameters was (−50,50) for *μ* and (0.1,100) for *σ*. Model comparison was performed based on the Akaike information criterion (AIC). Using other model comparison metrics such as the Bayesian information criterion (BIC) does not change the conclusions of our work.

In Figs. 3-4 and main text, we report the sum of the AIC difference relative to the best model across participants for each experiment, and report the 95% bootstrap confidence interval (95%-CI). We also computed two-sided paired t-tests based on the AICs obtained for each participant between the SEB and the TIM models. Likewise, we computed two-sided paired t-tests based on the guess rate parameter *g* obtained from each participant in the SEB model relative to the guess rates obtained in the TIM model. We report effect sizes as Cohen’s *d*. The qualitative predictions were computed based on the value with the highest density for each parameter at the population level. Each statistic was computed separately for each experiment and then averaged across experiments.

Details regarding the theoretical derivations of the SEB model and the analytical comparison between TIM and SEB models are given in detail in S1 Appendix–Supplementary Notes 1-5.

## Ethics statement

All data analysis from human participants is based on an openly available dataset [21], therefore no Institutional Review Board approval was required for this study.

## Supporting information

**S1 Appendix. Supplementary Figure, Tables and Notes**

## Acknowledgments

This work was supported by a European Research Council (ERC) starting grant (ENTRAINER) to R.P. This project has received funding from the European Research Council (ERC) under the European Union’s Horizon 2020 research and innovation programme (grant agreement No. 758604), and support from the U.S. National Science Foundation, under grant SES-DRMS-1949418 to M.W.

---

**Supplementary Table 1.**
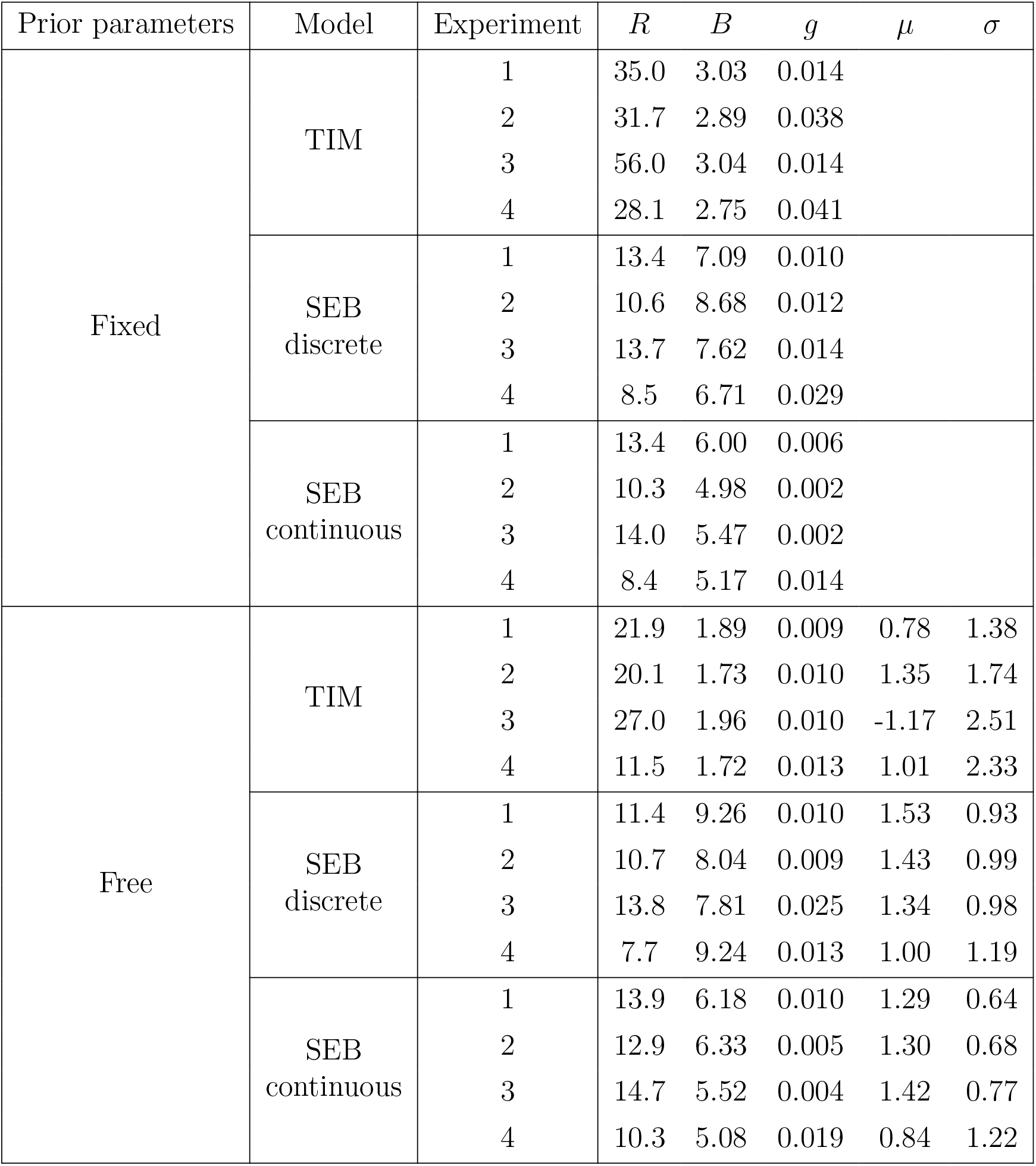
Parameter fits. Highest density of each parameter at the population level for each model and experiment.

| Prior parameters | Model | Experiment | $R$ | $B$ | $g$ | $\mu$ | $\sigma$ |
| --- | --- | --- | --- | --- | --- | --- | --- |
| Fixed | TIM | 1 | 35.0 | 3.03 | 0.014 |  |  |
|  |  | 2 | 31.7 | 2.89 | 0.038 |  |  |
|  |  | 3 | 56.0 | 3.04 | 0.014 |  |  |
|  |  | 4 | 28.1 | 2.75 | 0.041 |  |  |
|  | SEB<br>discrete | 1 | 13.4 | 7.09 | 0.010 |  |  |
|  |  | 2 | 10.6 | 8.68 | 0.012 |  |  |
|  |  | 3 | 13.7 | 7.62 | 0.014 |  |  |
|  |  | 4 | 8.5 | 6.71 | 0.029 |  |  |
|  | SEB<br>continuous | 1 | 13.4 | 6.00 | 0.006 |  |  |
|  |  | 2 | 10.3 | 4.98 | 0.002 |  |  |
|  |  | 3 | 14.0 | 5.47 | 0.002 |  |  |
|  |  | 4 | 8.4 | 5.17 | 0.014 |  |  |
| Free | TIM | 1 | 21.9 | 1.89 | 0.009 | 0.78 | 1.38 |
|  |  | 2 | 20.1 | 1.73 | 0.010 | 1.35 | 1.74 |
|  |  | 3 | 27.0 | 1.96 | 0.010 | -1.17 | 2.51 |
|  |  | 4 | 11.5 | 1.72 | 0.013 | 1.01 | 2.33 |
|  | SEB<br>discrete | 1 | 11.4 | 9.26 | 0.010 | 1.53 | 0.93 |
|  |  | 2 | 10.7 | 8.04 | 0.009 | 1.43 | 0.99 |
|  |  | 3 | 13.8 | 7.81 | 0.025 | 1.34 | 0.98 |
|  |  | 4 | 7.7 | 9.24 | 0.013 | 1.00 | 1.19 |
|  | SEB<br>continuous | 1 | 13.9 | 6.18 | 0.010 | 1.29 | 0.64 |
|  |  | 2 | 12.9 | 6.33 | 0.005 | 1.30 | 0.68 |
|  |  | 3 | 14.7 | 5.52 | 0.004 | 1.42 | 0.77 |
|  |  | 4 | 10.3 | 5.08 | 0.019 | 0.84 | 1.22 |

**Supplementary Table 2.** AIC of the model fit. The continuous version of SEB has the lowest AIC when the prior parameters are either fixed or free, which indicates a better fit to the behavioral data. In addition, the versions of the models with free prior parameters have lower AICs than the versions with fixed prior parameters. (Expt: Experiment).

| Prior parameters | Model | Expt. 1 | Expt. 2 | Expt. 3 | Expt. 4 | All Expts. |
| --- | --- | --- | --- | --- | --- | --- |
| Fixed | TIM | 55 136 | 63 016 | 56 126 | 68 453 | 242 731 |
|  | SEB discrete | 54 660 | 59 113 | 55 318 | 64 796 | 233 887 |
|  | SEB continuous | **53 663** | **57 732** | **53 810** | **63 347** | **228 552** |
| Free | TIM | 52 665 | 58 105 | 53 415 | 62 582 | 226 767 |
|  | SEB discrete | 52 536 | 55 954 | 52 427 | 62 247 | 223 165 |
|  | SEB continuous | **50 978** | **54 420** | **50 486** | **60 444** | **216 328** |

**Supplementary Figure 1.**
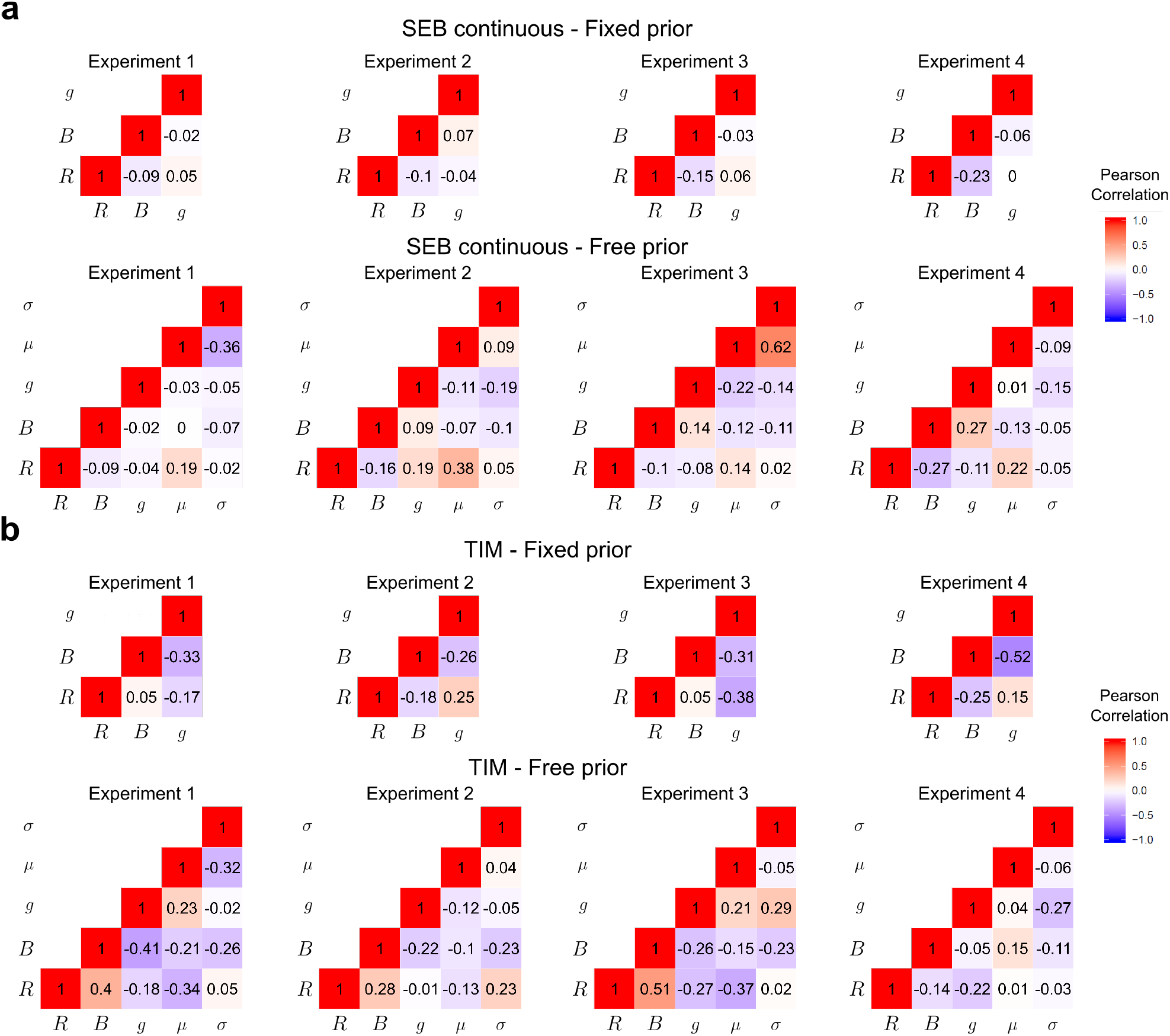
Correlations of parameter fits. **(a)** Pearson correlation of the parameter fits of the SEB continuous for each experiment with fixed (top) and free (bottom) prior parameters. **(b)** Pearson correlation of the parameter fits of the TIM model for each experiment with fixed (top) and free (bottom) prior parameters. The parameters across participants are weekly correlated suggesting that they are identifiable as confirmed in parameter recovery exercises.

## Supplementary Note 1. General specification and derivation of the logarithmic noisy encoding and Bayesian decoding model

The goal of this supplementary note is to introduce the noisy log-encoding Bayesian model that was elaborated in more detail elsewhere [2]. Additionally, we formalize the connection between the log-normal and power-law priors. The definitions and derivations specified here will serve as a basis to specify the information theoretical part of the model developed in Supplementary Note 2.

In this model, we assume that a stimulus of numerosity *n* generates an internal representation that is drawn from a distribution

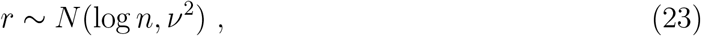

where the noise parameter *ν* is independent of *n*. Here we assume that the prior distribution from which the numerosity value *n* is drawn is given by a log-normal distribution

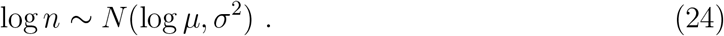

As stated in the main text, this distribution is qualitatively similar to the power-law distribution and also has many occurrences and applications in the statistics of human behavior. It is also generally present in various biological phenomena such as measures of length, area and weight of living organisms, and also present in neurophysiological observations such as distribution of firing rates across populations of neurons and intrinsic gain and synaptic weight in neural systems [3].

Based on these two assumptions (Eqs. 23 and 24), it follows that the distribution of log *n* conditional on the value of *r* will be a Gaussian distribution

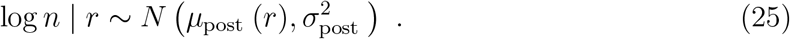

It follows that the conditional mean of log *n* is given by

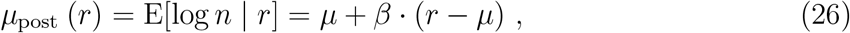

with the slope of this linear projection given by

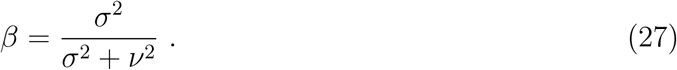

And the conditional variance is given by

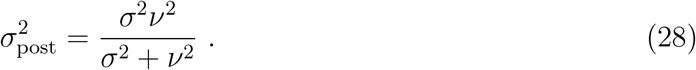

Here we consider the hypothesis that the participant’s numerosity estimate minimizes the MSE. Thus, the rule that is optimal under this objective is given where the estimate 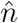 is defined by 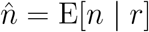 for all *r*. It follows from the properties of the log-normal distribution that the posterior mean is given by

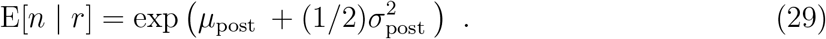

In this case, the Bayesian model predicts

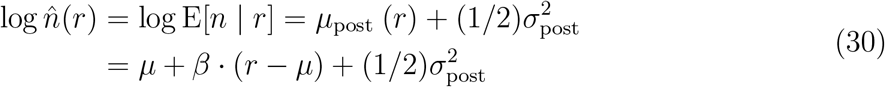

Given that *r* is a random variable, it follows that 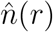 is also random variable. Thus, log 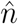 is normally distributed conditional on *n*

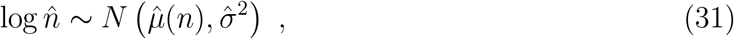

with the mean and variance of this conditional distribution given by

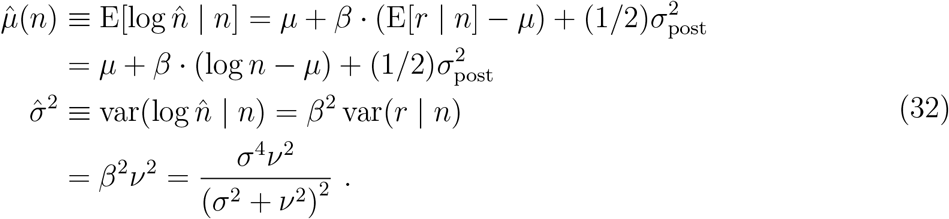

It then follows from the properties of the log-normal distribution that the expected value and variance of the numerosity estimators are given by

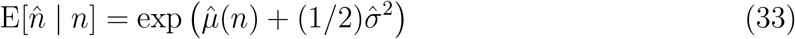

and

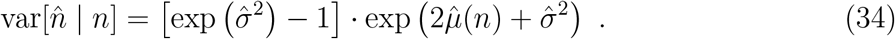

Finally, we can use the these equations to compute the ratio between the standard deviation and the expected value of the posterior estimators, i.e., the coefficient of variation (Eq. 14 in main text)

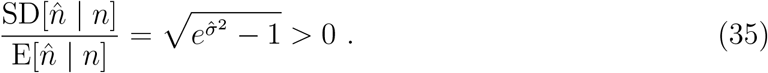

This expression does not depend on *n*, and therefore the log-encoding Bayesian model delivers the property of *scalar variability* discussed in the main text.

Note that in these calculations, only the *normalized prior* 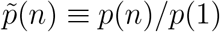 matters, and in fact the Bayesian posteriors can be well-defined even in the case of an improper prior (for which 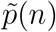 is well-defined, but there is no value for *p*(1) such that the implied density function *p*(*n*) will integrate to 1). All of the above calculations can be generalized to apply to any normalized prior of the form

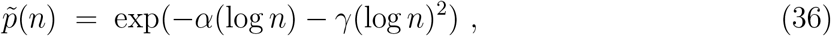

for some parameters *α, γ* with *γ ≥* 0. If *γ >* 0, this corresponds to a log-normal prior, with *μ* = (1 *- α*)*/*(2*γ*), *σ*^2^ = 1*/*(2*γ*). If instead *γ* = 0 but *α >* 0, this corresponds to an improper power-law prior, *p*(*n*) ∼ *n*^−*α*^.

In this latter case, the posterior implied by an internal representation *r* is again lognormal, as in equation (25), but now with parameters

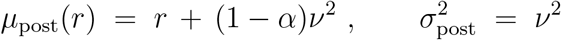

as limiting cases of equations (26) and (28). It then follows that the Bayesian posterior mean estimate 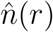 will be log-normally distributed conditional on the true value of *n*, as in equation (31), but with parameters

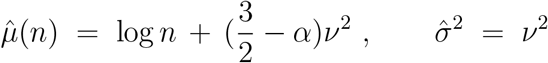

as limiting cases of the formulas given above. Hence the mean estimate will be given by

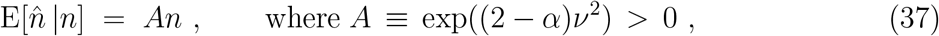

and the standard deviation of the estimates will again satisfy (Eq. 14), with the value of 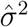 given above.

Thus even in the case of an improper prior of this kind, the optimal Bayesian estimate 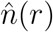 is well-defined, and we can derive the predicted distribution of 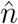 conditional on *n*, as a function of the model parameters. All priors in the family (Eq. 36) imply that the distribution of estimates should satisfy the property of scalar variability (Eq. 14). In the case of a log-normal prior (*γ >* 0), equation (33) implies that 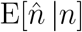 will be a strictly concave function of *n*, greater than *n* for all *n* below some critical value, and smaller than *n* for all *n* above the critical value. In the limiting case of a power-law prior (*γ* = 0), instead, equation implies that 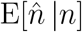 should be proportional to *n*, with either overestimation for all *n* (if *α <* 2) or underestimation for all *n* (if *α >* 2). In the special case of a power law with *α* = 2, the model implies that the optimal Bayesian estimate should be unbiased for all *n*.

## Supplementary Note 2. Logarithmic noisy encoding and Bayesian decoding under limited informational capacity and temporal sensory exposure

In this supplementary note we show that it is possible to formulate an efficient coding model of numerosity estimation as developed in Supplementary Note 1, but in which encoding precision depends on stimulus viewing time *t*.

Instead of assuming, as in Supplementary Note 1, that a stimulus of numerosity *n* results in an internal representation *r* that is a single draw from a probability distribution that depends on *n*, we suppose now that the internal representation *r* instead consists of the sample path of a Brownian motion *z*_*s*_ over a time interval 0 ≤ *s* ≤ *τ*, starting from an initial value *z*_0_ = 0. The drift *m* of the Brownian motion is assumed to depend on *n*, while its instantaneous variance *ω*^2^ is independent of *n*; the length of time *τ* for which the Brownian motion evolves is also independent of *n*, but depends on the viewing time *t*. In assuming sensory evidence given by a Brownian motion with a drift that depends on the stimulus, we follow a long modeling tradition that includes the popular drift-diffusion model [4]. Models of this kind have been used since Taylor, Lindsey and Forbes [5] to account quantitatively for the way in which the accuracy of perceptual judgments is affected by manipulations of viewing time.

More specifically, we assume that *m* is an affine transformation of the logarithm of *n*,

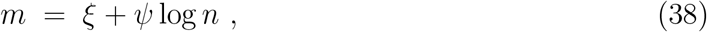

where the parameters *ξ* and *ψ* may depend of the statistics of a particular environment. We suppose that the choice of these coefficients is subject to a “power constraint” which requires the average value of *m*^2^ to be within some finite bound ^1^

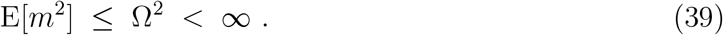

This bound on the amount of variation in the drift limits the precision with which different stimuli can be discriminated, for any given *τ*. The value of *τ* is assumed to grow linearly with the viewing time, up to some time bound *t*_*max*_,

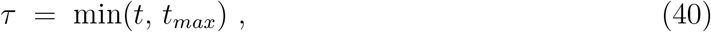

representing a constraint on the amount of evidence that can be acquired by the brain. The latter bound constrains the degree to which precision can be increased by further increases in viewing time, just as in the TIM model.

For any fixed value of *τ*, the final position *z*_*τ*_ of the Brownian motion at time *τ* is a sufficient statistic for the information contained in the sample path about the value of *n*. Hence Bayesian decoding of the information contained in the sample path will yield the same result as if the internal representation is assumed simply to be the scalar random variable *z*_*τ*_, with distribution

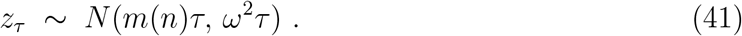

Alternatively, we may suppose that the internal representation of *n* is given by the scalar random variable *r* ≡ *z*_*τ*_ */τ*, which contains the same information as the variable *z*_*τ*_. Under this representation of the sensory evidence, *r* is a draw from a distribution

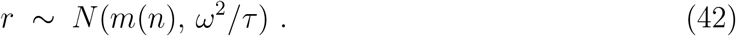

Equation (42) effectively states that *r* is the output of a *Gaussian channel* with input *m* [1]; hence the problem of optimally choosing the function *m*(*n*) is equivalent to an *optimal encoding* problem for a Gaussian channel. The capacity *C* of such a channel is a quantitative upper bound on the amount of information that can be transmitted regardless of the encoding rule, which is equal to

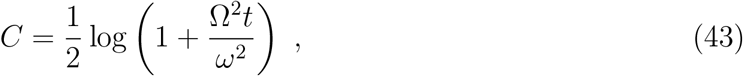

an increasing function of Ω*/ω* as well as of *t*. Here we suppose that the goal is to design a system that minimizes the mean squared error of the estimate 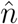 when *n* is drawn from a log-normal prior distribution (Eq. 24).

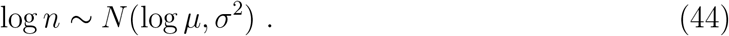

Note that the estimate 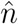 depends on *r*_*n*_. We re-express *r*_*n*_ as a function of the transformed variable 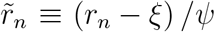, thus we can equivalently treat 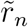 as the internal representation, and it follows that 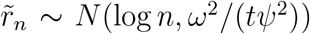. Following the definitions provided in Supplementary Note 1, it follows that we have a noisy log-encoding with variance *ν*^2^ = *ω*^2^*/*(*tψ*^2^). It then follows that the MSE in the case of any encoding rule is given by Eq. 30, and the MSE associated with this rule will be given by

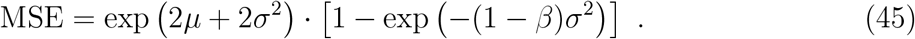

Recall that *β* is a decreasing function of *ν* (Eq. 27), and therefore in order to make the MSE as small as possible it is desirible to make *ν* as small as possible. Given that *ν*^2^ = *ω*^2^*/*(*tψ*^2^) is follows that we would like to make *ψ* as large as possible, consistent with the power constraint in Eq. 39. Thus, for the case of the log-normal prior the power constraint becomes

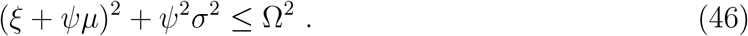

The maximum value of *ψ* consistent with this constraint is achieved when

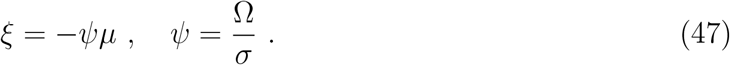

In this case the encoding noise is given by

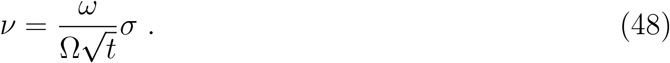

Defining *R* ≡ Ω*/ω*, we can define the encoding noise of numerosity estimation

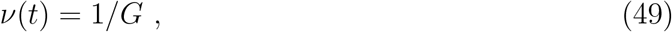

where

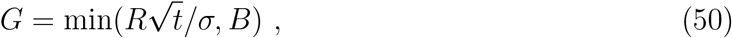

with *B* a maximum biologically allowed bound on sensory precision (similar to the assumption of the TIM model).

The precision of numerosity encoding is given by *ν*(*t*) = 1*/G*, where 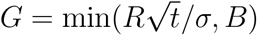 and *B* a maximum biologically allowed bound on sensory precision related to 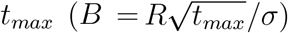.

## Supplementary Note 3. Recapitulation of the Gaussian channel capacity derivation

For convenience to the reader, the goal of this supplementary note is to recapitulate the derivation of the Gaussian channel capacity presented in Cover and Thomas [1] based on the notation used in our work, thus clarifying the connection to the solution of our SEB model under capacity constraints derived in Supplementary Note 2.

Suppose that *Y* is the output of a channel with input *X* + *Z*, where *X* is the signal and *Z* the noise. We assume the noise is drawn from a Gaussian distribution with variance *ω*^2^*/t* and mean 0.

The goal is to find the maximum achievable channel capacity *C* by maximizing the mutual information *I*(*X*; *Y*) for a given power constraint Ω^2^

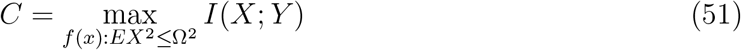

It can be shown that

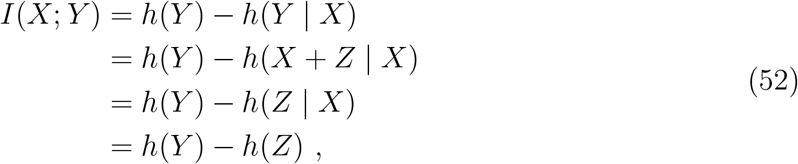

where in general *h*(*X*) is defined as the entropy of the channel *X*. Here, we will use two results. First, the entropy of the input of a Gaussian channel, for a given noise *Z* with variance *ω*^2^, is given by

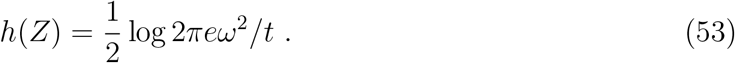

Second,

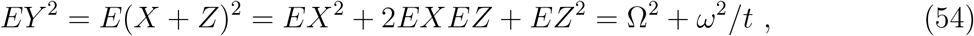

given that *X* and *Z* are independent and *EZ* = 0. This means that the entropy of *Y* is bounded by

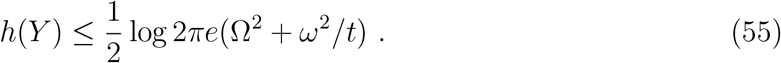

Thus, replacing Eqs. 53 and 55 in Eq. 52 gives

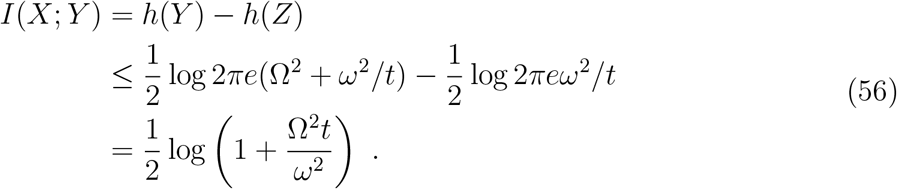

## Supplementary Note 4. Influence of the prior on the crossover point

The crossover point corresponds to the numerosity at which participants switch from over-estimation to underestimation. Here, we use the analytical solutions of the SEB model to show how the crossover point depends on elements of the prior distribution and the capacity constraints. More specifically, we can find the numerosity *n*^∗^ at which the crossover will occur. In other words we can find the numerosity at which the following equality holds

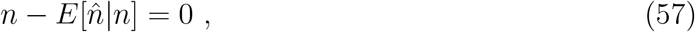

or alternatively

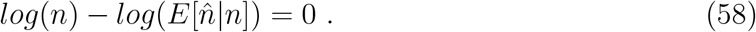

Using the solutions in Eqs. 25, 26 and 31 it is possible to show that

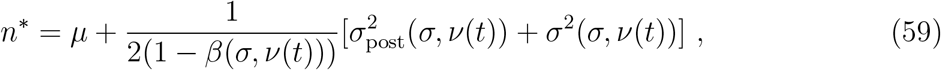

where recall that *μ* is the prior location, *σ* is the prior spread, and *ν*(*t*) is the endogenized encoding noise function for a given time *t* (and also for a given encoding capacity). Note that none of the elements of expression in the second term in the right-hand side of the equality depend on the prior location *μ*, which means that changes in *μ* without changes prior spread or noisy coding elements will induce shifts in the direction of the prior location. This also means, and while not obvious from this expression, that positive changes in *σ* will induce a positive shift of the crossover point.

## Supplementary Note 5. Comparison between the family TIM and SEB models

In this supplementary note, we compare the kind of information theoretical model that does not incorporate Bayesian inference (the TIM model defined in the main text), and the general family of Bayesian encoding-decoding models. This illustrative comparison is not directly applicable to our numerosity estimations, but can be solved analytically and is instructive. Nevertheless, we provide numerical simulations that are applicable to numerosity estimation in the main text which confirm the main predictions presented in this note (see Fig. 2 in main text).

### Illustrative comparison between the family of TIM and SEB models

In this illustrative example, the two models can be solved analytically, thus allowing to highlight the commonalities and differences between both models in an intuitive manner.

The TIM model proposes a method to infer how the distribution of estimates 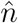 should vary depending on the true stimulus magnitude *n*. The goal is to find the response distribution 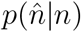 that minimizes the mean squared error (MSE)

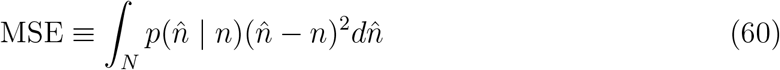

subject to the constraint that (Eq. 18 in main text)

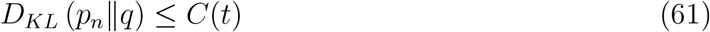

where *p*_*n*_ is the distribution of possible responses conditional on *n, q* is the “prior” distribution, *D*_*KL*_ (*p*∥ *q*) is the Kullback-Leibler divergence, and *C*(*t*) is a positive bound that depends on the amount of time *t* for which the stimulus is presented.

The TIM model can be developed further by specifying that *q* is given by the prior distribution from which *n* is expected to be drawn, and that *C*(*t*) increases linearly with time, i.e., that *C*(*t*) = *c*· *t* for some *c >* 0, up to some finite bound B. Assuming that the prior is known, the model thus has only a single free parameter (the value of *c*) to predict the distribution of responses, as a function of both *n* and *t*, for all values of *t* below some upper bound.

We compare the predictions of this kind of model to the alternative Bayesian model, according to which (i) estimates are based on a noisy internal representation *r* of the stimulus magnitude *n*, which consists of a sequence of independent draws of a signal, the distribution of which depends on *n*, and with the number of draws in the sequence growing with *t*; and (ii) given the noisy internal representation, the participant’s estimate is given by 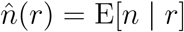: Note that the computation of this last conditional expectation must be relative to a particular prior distribution from which *n* is expected to be drawn. Given the distribution of possible samples *r* for each *n* and *t*, we can use the assumed response rule to derive a predicted distribution of responses 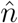 for any specification of (*n, t*).

We make the Bayesian model example more specific by assuming that *r* is the cumulative value at time *t* of a Brownian motion that starts from the initial value *r* = 0, with a drift *m*(*r*) that depends on the stimulus and an instantaneous variance *ω*^2^ that is independent of the stimulus. If we further assume that *m*(*n*) = *μ*· *n* for some *μ >* 0; then the model’s predictions depend only on a single parameter, the value of *γ* ≡ *μ/ω*; again assuming that the prior is known. We thus have two one-parameter models, each of which makes precise predictions for the distribution 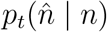 for any *n* and *t*. Thus, in each model, the single free parameter determines how rapidly the precision of estimates should improve with increasing viewing time.

In this example, we suppose that the prior distribution for *n* is Gaussian, and let it be given by *N* (0, *σ*^2^). Here, we economize in notation by assuming that the prior mean is zero; the formulas that follow hold regardless of this, but *n* should be understood as the stimulus magnitude relative to the prior mean, and 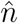 as the response relative to the prior mean.

In the case of the Bayesian model, the information contained in the noisy internal representation is equivalent to that for a model in which the available information is a noisy measurement, *r* ∼ *N* (*n*, (*γ*^2^*t*)^−1^), the precision of which grows linearly with *t*. The optimal Bayesian estimate is then 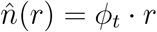, where

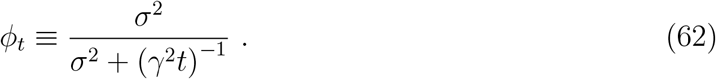

From this, it follows that the conditional distribution of responses for any time *t* will be given by

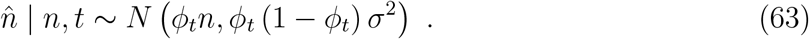

For the TIM model instead: the first order conditions for minimization of Eq. 19 subject to Eq. 18 require that

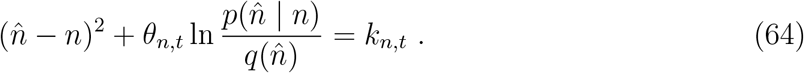

for all 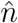, where *θ*_*n,t*_ is the Lagrange multiplier associated with the capacity constraint given in Eq. 18 for given choices of *n* and *t*, and *k*_*n,t*_ is a constant of integration. This equation can be solved for 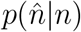for each 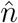, given values for *k*_*n,t*_ and *θ*_*n,t*_. We choose *k*_*n,t*_ so as to imply a PDF 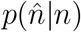 that integrates to 1. We see that the resulting distribution for 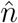 is Gaussian

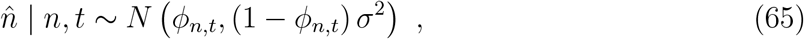

where the bias coefficient *ϕ*_*n,t*_ corresponds to

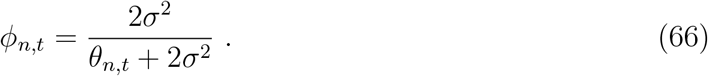

The value of *θ*_*n,t*_ is chosen so as to imply that the constraint given in Eq. 18 holds with equality. Computing the KL divergence, we see that this holds if and only if

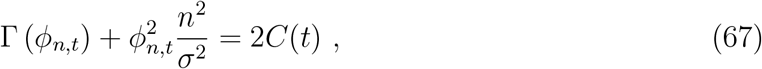

where

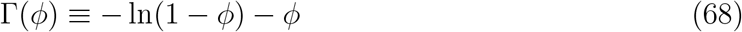

for any 0 *< ϕ <* 1.

For any *n*, we note that Γ(*ϕ*) is a continuous, monotonically increasing function of *ϕ*, approaching 0 as *ϕ* → 0 and becoming unboundedly large as *ϕ* → 1. Hence, for any *n* and any *C*(*t*) *>* 0, equation 67 has a unique solution satisfying 0 *< ϕ*_*n,t*_ *<* 1. We further observe that for a fixed value of *n*, increasing *C*(*t*) increases the value of *ϕ*_*n,t*_; and for a fixed *C*(*t*), increasing the value of |*n*| increases the value of *ϕ*_*n,t*_.

Based on these analytical solutions, these results reveal some important similarities between the predictions of the TIM and the Bayesian models: for large enough *t* and allowing *C*(*t*) to grow as function of *t*, then both models imply

1. 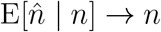, and
2. 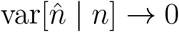.

Nonetheless, there are also several notable differences in the predictions of the two models. Here we provide a detailed explanation of these differences which were already mentioned in the main text:

1. *The quantitative dependence of estimation bias on viewing time*. While both models predict that *ϕ*_*n,t*_ should increase from 0 (for *t* = 0) to 1 (as *t* → ∞), they do not imply the same rate of increase in *ϕ*_*n,t*_ as *t* increases. The Bayesian model implies that

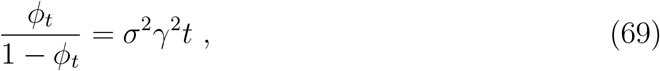

for any *n*. Hence for small *t, ϕ*_*t*_ ∼ *t*, while for large *t*, (1 −*ϕ*)^−1^ ∼ *t*. Instead, if in the TIM model we assume that *C*(*t*) = *c* ·*t* for all *t*, then for any *n* ≠ 0, one can show that the solution to Eq. 67 satisfies *ϕ*_*n,t*_ *t*∼ ^1*/*2^ for small *t*, while (1− *ϕ*)^−1^ ∼ *e*^2*ct*^ for large *t*. Thus regardless of the parameters *γ* and *c* for the two models, we see that the TIM model implies faster growth of *ϕ*_*n,t*_ as *t* increases, both for sufficiently small values of *t* and for sufficiently large values of *t*.
2. *The relationship between estimation bias and the variability of estimates*. Again fixing some single value of *n* ≠ 0, and considering the implied distribution of estimates for different viewing times, we see that the two models do not imply that 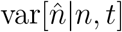 covaries with the bias in the same way. The TIM model implies that the variance falls monotonically with increases in *ϕ*_*n,t*_ (and hence that the variance falls monotonically with time, for any *n*). The Bayesian model instead implies that increases in *ϕ*_*t*_ first increase variance (while *ϕ* remains below 1/2), and then reduce variance again (once *ϕ*_*t*_ *>* 1*/*2). The difference in predictions is especially stark in the case of small viewing time. As *t* → 0, the Bayesian model’s variance falls to zero (estimates are equal to the expected value of the prior), while the TIM model implies that this is the case in which estimates should be most variable (estimates are simply samples from the prior distribution, regardless of the value of *n*).

There are other differences between the two models that we do not highlight here as they are not strictly relevant to the discussion of this article.

Taken together, we developed an example in which analytical analyses allowed us to examine commonalities and differences between the two models. While the exact predictions of these differences do not hold for the specific application of the numerosity estimation models developed for the TIM model and the noisy log-encoding Bayesian model (see Supplementary Notes 1 and 2), these general differences make the two numerosity models identifiable, and thus generate different qualitative predictions. In particular, the two differences highlighted above cause the TIM model not to provide a general account of the scalar variability principle. That is, the ratio between variability and expected value estimations grows more rapidly in the TIM model relative to the log-encoding Bayesian model.

## Footnotes

1 The rationale behind the choice of the squared *m* is the following: The signals can take any value over the real line. If we assume that the status quo of no-message (or no energy being spent) is 0, then any deviations from 0 that lead to decodable information should be considered including negative values leading to energy expenditure.

